# LNGCN: a distance-aware continuous-time graph framework for PPI prioritization in experimental screening

**DOI:** 10.64898/2026.04.30.721835

**Authors:** Yueming Xiao, Yifan Zheng, Yu Hua, Jiahua Peng, Jinliang Liu, Yuan Qu, Jizhuang Xu, Rao Fu, Qiuting Qian, Martin Kosar, Yuehai Ke, Ying Chi

## Abstract

High-throughput PPI screening requires not only interaction discrimination but also stable, calibrated scores for candidate prioritization and efficient experimental allocation. Conventional graph neural networks may over-smooth deep residue representations, reducing score separation among high-confidence candidates. We therefore developed LNGCN, a distance-aware continuous-time graph framework for experimental PPI prioritization. LNGCN integrates residue-level structural graphs with liquid neural dynamics, uses radial distance to drive continuous graph evolution, and applies hierarchical calibration to convert raw outputs into prioritization scores. It performed robustly on balanced human, 1:10 imbalanced, and cross-species yeast benchmarks. On the imbalanced dataset, the top 5% of candidates achieved 85.1% precision. LNGCN also preserved residue-level representation diversity better than conventional GCN variants. In FGF23-FGFR1c-*α*-Klotho, SHP2 signaling, Tdk1 oligomer-dependent binding, and a TPR study, it enriched known partners near the ranking top, and experimentally supported novel interactions. Overall, LNGCN enables sustained enrichment of experimentally relevant PPI candidates.

## INTRODUCTION

Protein-protein interactions (PPIs) underpin cellular signaling, molecular complex assembly, and gene-expression regulation^1^. Although experimental assays remain indispensable for identifying PPIs, their cost, limited throughput, and technical variability restrict validation across large candidate spaces. In practical screening, a computational model must do more than classify protein pairs as interacting or non-interacting. It should assign stable continuous scores to thousands of candidates so that the most promising pairs can be selected when experimental capacity is limited. The usefulness of PPI prediction therefore depends not only on classification accuracy, but also on whether true interactions are enriched near the top of a ranked list.

Current PPI models integrate protein sequences, evolutionary profiles, pretrained protein language models, and three-dimensional structures. Sequence-based and language-model approaches capture contextual and evolutionary patterns from large protein corpora^2^, whereas structure-aware methods represent proteins as residue-level graphs and use graph convolutional networks (GCNs) to encode spatial organization^3,4^. GCNs have been applied to interface-residue identification^5^, PPI prediction^6^, protein–ligand modeling^7^, and edge-aware structural learning^8^. However, most studies still formulate PPI prediction primarily as binary classification and emphasize metrics such as AUROC. These metrics measure overall discrimination but do not directly answer a practical question: how many true interactions can be recovered when only a small fraction of the highest-ranked candidates can be tested?

This limitation is especially important in class-imbalanced pools, where negative pairs greatly outnumber positives. Under such conditions, high accuracy or AUROC does not necessarily translate into effective early enrichment^9^. PPIs prioritization should therefore also be evaluated using Precision@k, Recall@k, normalized discounted cumulative gain (NDCG), and cumulative recovery analyses^10,11^. Cross-species evaluation is likewise important because strong within-species performance may partly reflect species-specific signatures rather than transferable interaction-related patterns^12^.

Graph representation may further affect ranking quality. Conventional message passing propagates information through discrete neighborhoods, whereas residue-specific spatial positions are rarely incorporated as explicit dynamic signals. Repeated aggregation can make node embeddings increasingly similar with increasing depth, a phenomenon known as over-smoothing^13–15^. Importantly, such information loss exerts distinct effects on binary classification and candidate prioritization. Binary classification merely aims to separate positive from negative pairs, while prioritization further demands preservation of subtle and consistent score differences across high-confidence candidates. Excessive homogenization may therefore weaken distinctions among structural regions and compress the score range at the top of the candidate list. In parallel, raw neural-network outputs are often overconfident and cannot be interpreted directly as reliable screening scores. A practical framework must both preserve representational diversity and transform raw predictions into empirically interpretable prioritization values. These values should be distinguished from binding free energies^16^. Its objective is to increase the enrichment of true interactions within a limited set selected for experimental validation, rather than to estimate the physical strength of binding.

Continuous-time neural dynamics provide a strategy for addressing the representation problem. Liquid neural networks (LNNs) and closed-form continuous-time networks (CfCs) describe hidden-state evolution through learnable dynamics and adaptive decay^17 18^. Rather than relying solely on stacked discrete graph layers, continuous dynamics allow residue-specific spatial information to influence state evolution directly. We therefore combine radial distances with graph-neighborhood aggregation as driving signals in an ordinary differential equation, enabling position-dependent residue trajectories while retaining local structural context. To address the second problem, post hoc calibration is applied to reduce overconfidence and convert the resulting raw outputs into empirical priority scores suitable for candidate screening.

Based on this design, we developed LNGCN, a distance-aware continuous-time graph framework for experimental PPI screening prioritization. A CfC module compressed residue-level features while preserving radial-distance information. Multiscale graph convolution was embedded within an LNN-based dynamical system so that structural neighborhoods and spatial position jointly guide residue-state evolution. Protein representations were combined through symmetric pair fusion, ensuring invariance to input order, and raw outputs were calibrated into empirical priority scores. This score represents the tendency of the model to predict the relative priority of interactions, rather than the strict probability of biological physical binding.

We evaluated LNGCN across discrimination, ranking, transferability, representation preservation, and biological utility. Performance was assessed on balanced human data and a 1:10 imbalanced setting using Top-k, NDCG, enrichment, and recovery metrics. Cross-species transfer was examined on yeast, while residue-level analyse assessed node variance, spatial separation, and effective embedding rank. Case studies involving the FGF23–FGFR1c–*α*-Klotho complex^19^, SHP2 signaling, and Tdk1 oligomeric states^20,71^ evaluated mechanism-consistent ranking, and perturbation analyses examine structurally plausible score-sensitive regions. Finally, a TPR experimental application prioritized partners, including Co-IP support for ELAVL1-TPR and RALY-TPR. Together, these results prove that LNGCN can translate PPI predictions into candidate prioritization with practical guidance value.

## RESULTS

### Overview of the LNGCN protocol

LNGCN is a distance-aware continuous-time graph framework designed for high-throughput PPI candidate screening. It aims to generate stable, continuous, and calibrated interaction-priority scores within large candidate spaces containing both potentially interacting and non-interacting protein pairs. These scores support efficient candidate selection when experimental resources are limited. The model follows a Siamese architecture, in which two input proteins are encoded through shared modules before their graph-level representations are combined for pairwise interaction prediction (Figure 1). This design enables independent protein encoding while preserving symmetry at the pair level.

**Figure 1:**
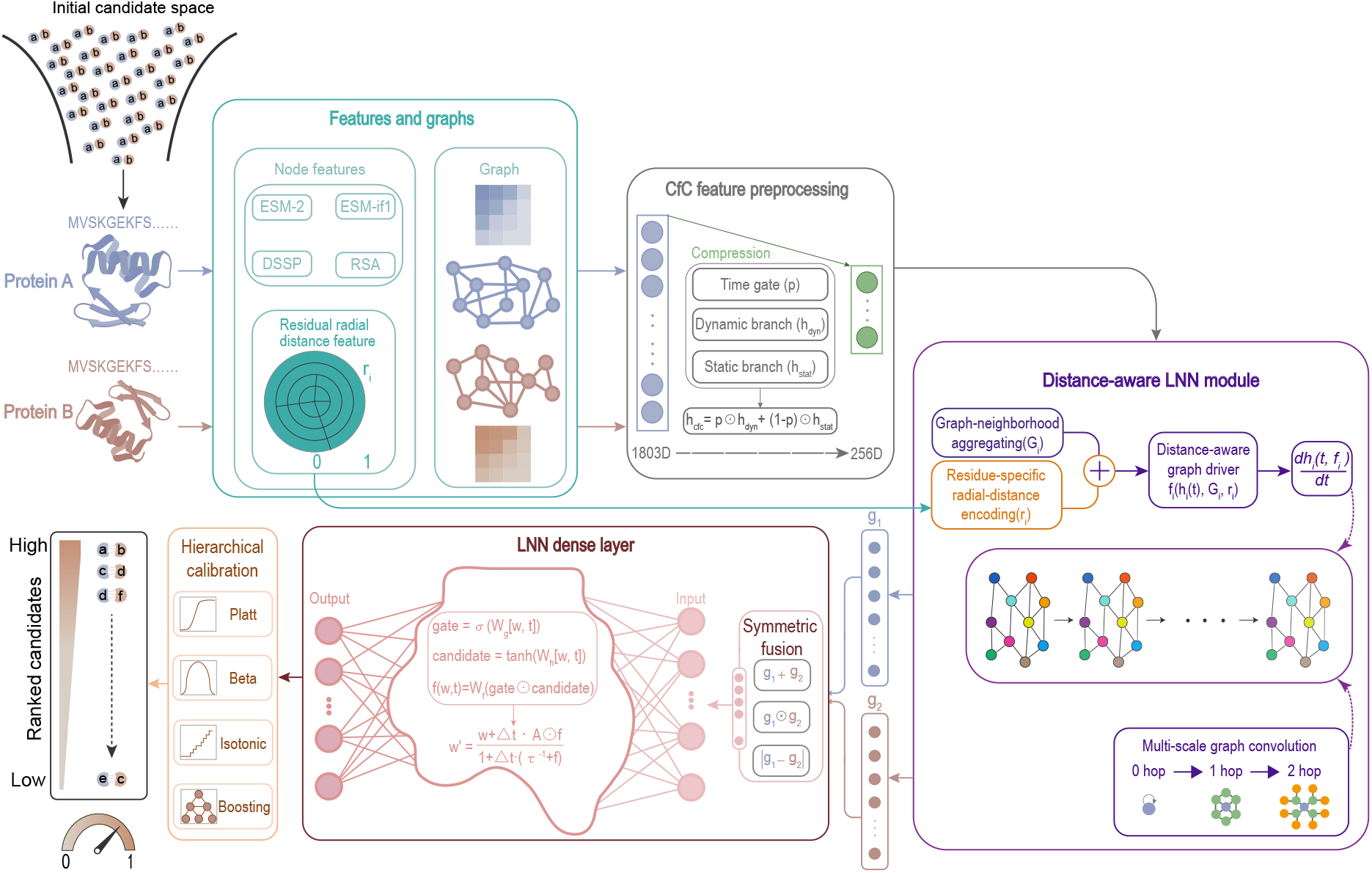
Overview of the LNGCN framework for distance-aware continuous-time PPI candidate prioritization. Candidate protein pairs are first represented as residue-level structural graphs integrating multimodal sequence and structural features, including ESM-2^22^, ESM-IF1^47^, DSSP^24^, RSA^25–28^, and residue radial distance feature. CfC-based preprocessing compresses the high-dimensional residue features through gated dynamic and static branches while retaining radial distance as an explicit spatial signal. The distance-aware LNN module then combines multi-scale graph-neighborhood aggregation with residue-specific radial-distance encoding to drive continuous residue-state evolution. The resulting protein representations are processed by an LNN dense layer and combined through symmetric fusion to generate an order-invariant raw interaction score. Finally, Platt, Beta, Isotonic, and Boosting calibration methods are hierarchically integrated to transform the raw output into an empirically calibrated prioritization score. These scores are used to rank candidate protein pairs for PPI screening.

The protocol integrates multimodal residue features, graph convolutional networks^4^, liquid neural networks^17^, and closed-form continuous-time networks^18^. It contains five stages: (1) residue-level structural graph construction and extraction of high-dimensional static features together with residue radial distance; (2) CfC-based feature compression while retaining radial distance as an explicit spatial signal; (3) distance-aware continuous graph evolution through a graph-driven LNN module; (4) symmetric protein-pair fusion and LNN dense classification; and (5) multi-method probability calibration that converts raw outputs into prioritization scores. Thus, LNGCN translates protein sequence and structural information into calibrated PPI probabilities for candidate ranking in screening workflows.

### Datasets

For model training, high-confidence human protein-protein interactions (PPIs) with STRING combined scores greater than 0.9 were selected as positive samples^29^. Negative samples are generated by randomly matching from all the known protein interaction pairs to create protein pairs that do not exist in the interactions. Protein sequences and AlphaFold-predicted structures^30^ were obtained from UniRef^31^. The main dataset was divided into training, validation and independent test sets, as summarized in Table 1.

**Table 1:**
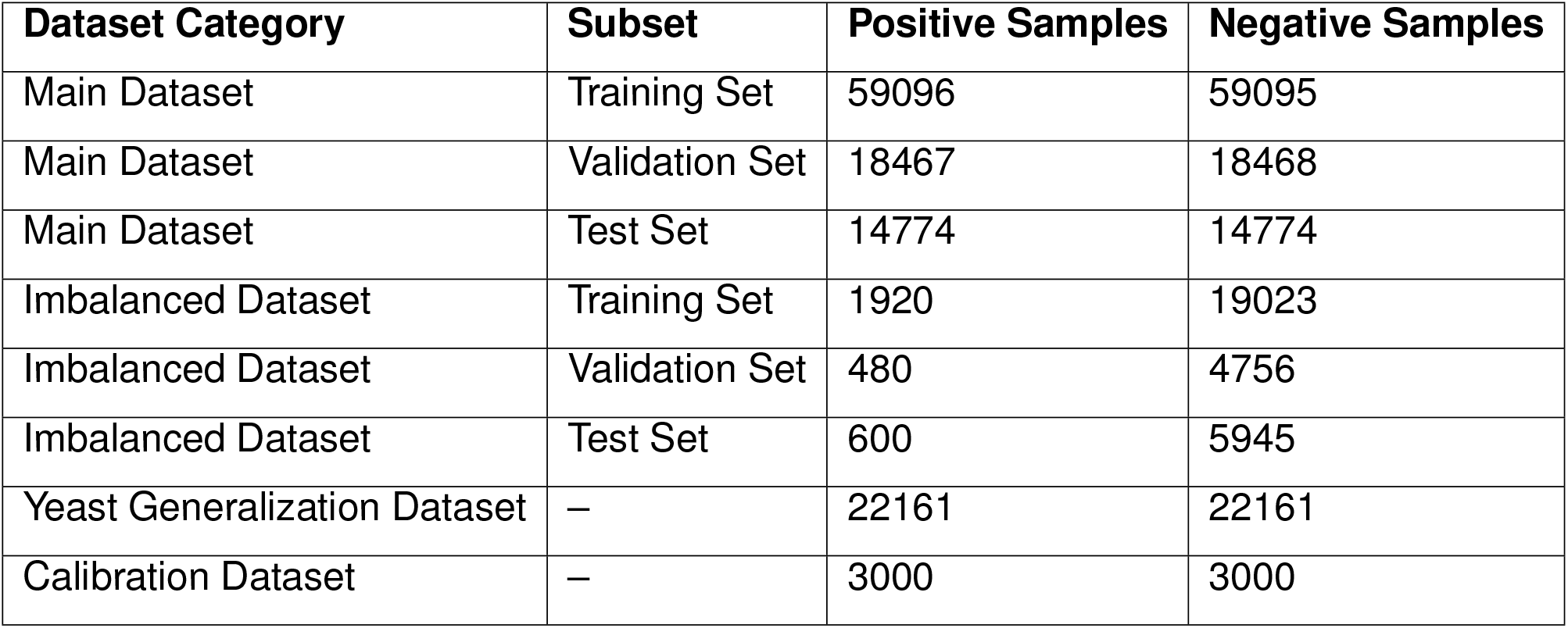
Sample Size Distribution of Datasets.

To evaluate model performance under a more realistic class-imbalanced setting, we additionally used an independent human PPI dataset derived from Zhang et al.^32^. The positive set contained 3,000 high-confidence PPIs shared across STRING^29^, UniRef^31^, and BioGRID^33^. For negative sampling, 30,000 human protein pairs without recorded interaction evidence were initially generated. Pairs containing UniProt entries flagged as obsolete, deleted, or subject to major revision were removed, resulting in 29,724 putative negative pairs and an approximate positive-to-negative ratio of 1:10. This dataset was evaluated separately from the primary benchmark to assess candidate prioritization under substantial class imbalance.

Cross-species generalization was assessed using an independent yeast PPI dataset obtained from STRING^29^. For calibration, we used 3000 interactions from Zhang et al.^32^ as positives and 3000 negatives (2000 random non-interactions plus 1000 from Negatome^35^). The calibration set was not used for training the base classifier or evaluating independent test discrimination. This calibration aligns model output with true biophysical interaction probabilities, reducing false-negative interference. For negative samples, we combined 2000 random non-interactions with 1000 experimentally validated non-interactions from Negatome^35^. The data employed by this calibration mechanism enables model outputs with true biophysical interaction probabilities, reducing false-negative interference.

We adopted multiple metrics to benchmark basic classification performance: Area Under the Precision-Recall Curve (AUPRC), AUROC, accuracy, precision, recall, F1-score, and Matthews Correlation Coefficient (MCC). Receiver operating characteristic (ROC) and precision-recall (PR) curves were also plotted. For ranking performance, we further used Precision@k, Recall@k, and normalized discounted cumulative gain (NDCG@k). Precision@k represented the proportion of true interactions among the top-k ranked candidates, whereas Recall@k measured the proportion of all true interactions recovered within this candidate subset. NDCG@k additionally considered the specific positions of true interactions within the top k candidates, with k=50, 100, and 500. Correct candidates appearing closer to the top of the ranked list were assigned greater weight.

### LNGCN establishes a reliable predictive foundation for PPI prioritization

On the STRING^29^ human benchmark, five-fold cross-validation yielded stable performance. LNGCN achieved a mean AUPRC of 0.9806 ± 0.0046 and a mean AUROC of 0.9805 ± 0.0051 on the independent test folds (Figure 2a), together with an F1 score of 93.93% and an MCC of 0.8787. Cross-fold variation was small, supporting the robustness of the model outputs. Fold-level results are provided in Supplemental information S1.

**Figure 2:**
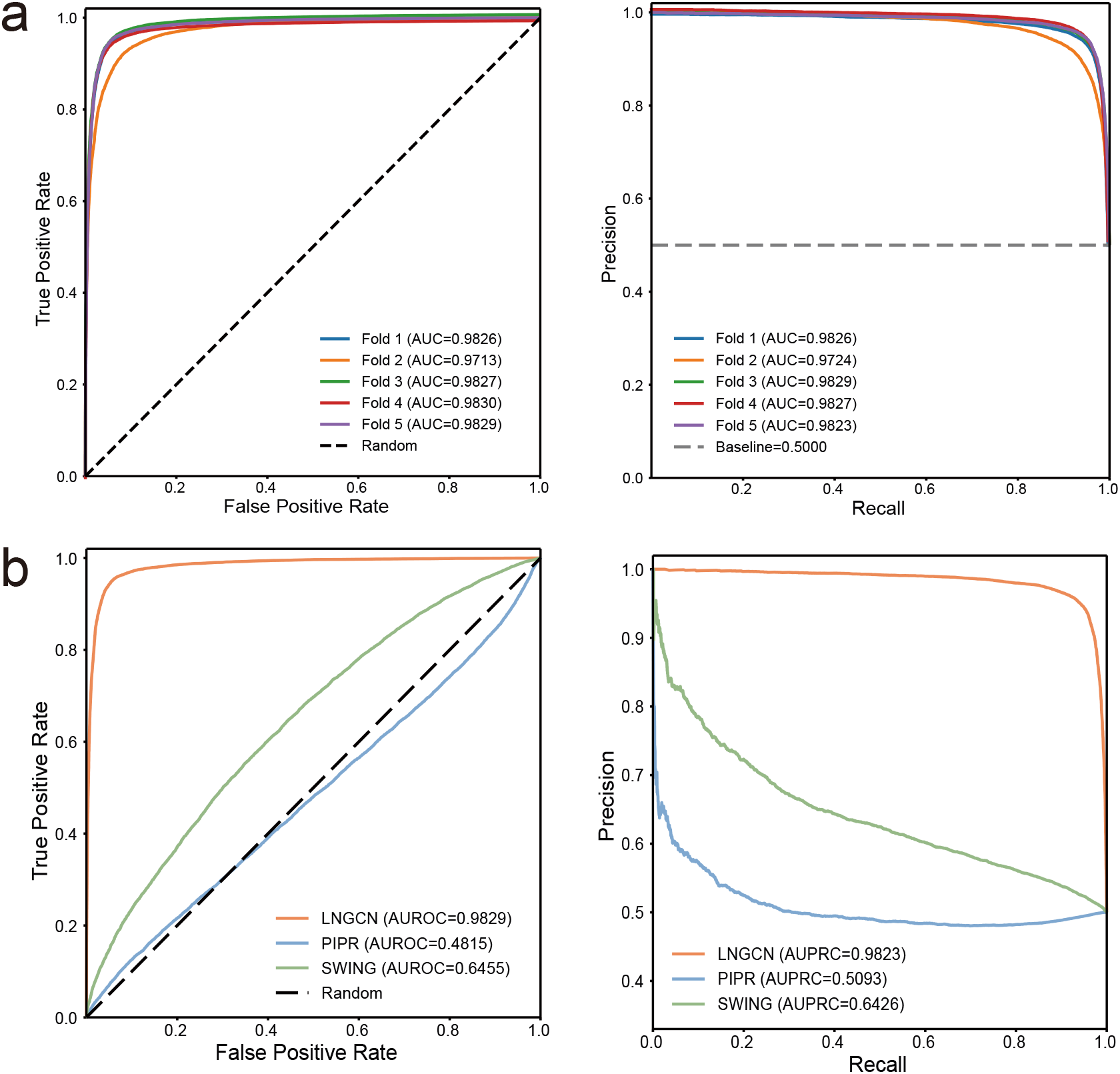
Model performance evaluation on balance dataset. (A) Benchmark ROC and PR curves. (B) ROC curve comparison between our model and two existing methods on the balanced dataset.

We further compared LNGCN with PIPR^36^, a Siamese residual recurrent convolutional model, and SWING^2^, a sliding-window interaction language model. Under the same benchmark setting, LNGCN outperformed both baselines across all major metrics (Table 2, Figure 2b). In fold 5, LNGCN retained higher overall performance, whereas PIPR and SWING showed lower AU-ROC and MCC values. These findings support the contribution of residue-level structural graphs and continuous-time representation learning, while providing stable continuous scores for downstream PPI prioritization in class-imbalanced candidate spaces.

**Table 2:** Performance comparison under the same benchmark test dataset.

| Method | AUPRC | AUROC | Accuracy | Precision | Recall | F1-Score | MCC |
| --- | --- | --- | --- | --- | --- | --- | --- |
| LNGCN | **0.9823** | **0.9829** | **0.9468** | **0.9417** | **0.9525** | **0.9471** | **0.8936** |
| PIPR | 0.5091 | 0.4815 | 0.4720 | 0.4840 | 0.8465 | 0.6159 | −0.0845 |
| SWING | 0.6426 | 0.6455 | 0.6010 | 0.5937 | 0.6400 | 0.6160 | 0.2027 |

### Top-ranked recovery under imbalanced screening demonstrates prioritization utility

To assess LNGCN under class imbalance, we evaluated it on the 1:10 human PPI dataset of Zhang et al.^32^. This class distribution more closely reflected real screening scenarios, in which true positive interactions were sparse within a large candidate pool. LNGCN outperformed RF2-PPI across classification metrics (Figure 3a). Across five test folds, mean AUPRC, AUROC, and accuracy were 0.7522, 0.9427, and 0.9390, respectively (Figure 3b). LNGCN retained strong discrimination despite the more challenging class distribution.

**Figure 3:**
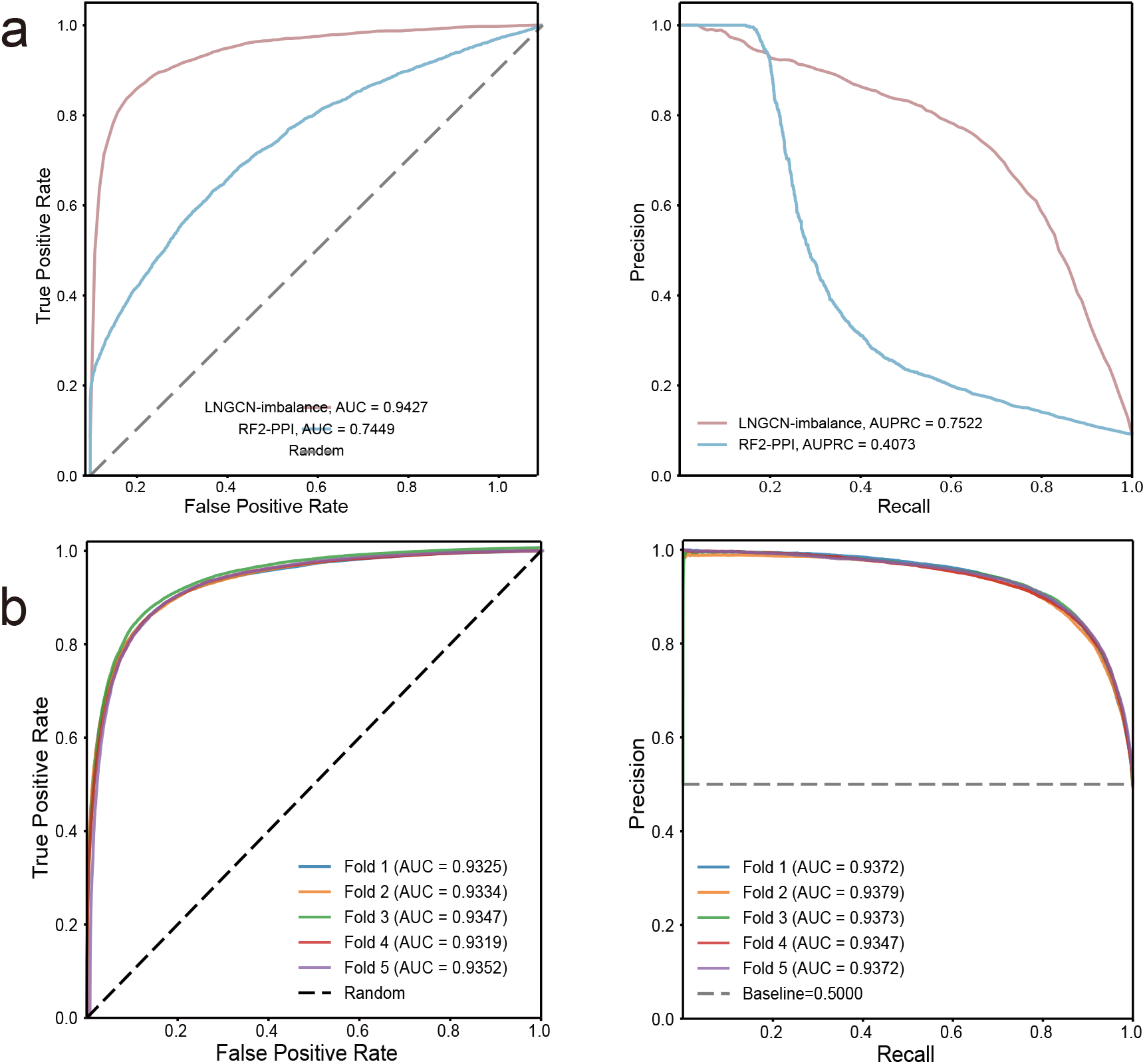
Classification performance of LNGCN on the 1:10 imbalanced human PPI benchmark. (A) Receiver operating characteristic ROC and PR curves comparing LNGCN with RF2-PPI under the same benchmark setting. (B) ROC and PR curves obtained from five-fold cross-validation of LNGCN on the imbalanced dataset. Each colored curve represents the performance on an independent test fold.

LNGCN also achieved consistent early enrichment. Precision@50 and NDCG@50 reached 0.9620 ± 0.0426 and 0.9637 ± 0.0400, respectively, while Precision@100 remained 0.9350±0.0460. These cutoffs represented only 0.76% and 1.53% of the candidate space. At the top-500 cutoff, LNGCN achieved a precision of 0.7444 ± 0.0190, recall of 0.6203 ± 0.0159, and NDCG of 0.7772 ± 0.0166. In other words, screening approximately 7.64% of candidates recovered an average of 62.0% of all true interactions.

Proportion-based analysis further highlighted this enrichment (Figure 4a-b). Within the top 5% of the ranked list, precision and NDCG were 0.8445 ± 0.0350 and 0.8640 ± 0.0307, respectively. Approximately 277 of the 328 prioritized pairs per fold were true interactions, compared with approximately 30 expected under random sampling, corresponding to a 9.2-fold enrichment. When the screening range was expanded to the top 20%, LNGCN recovered 87.2% of all positive samples.

**Figure 4:**
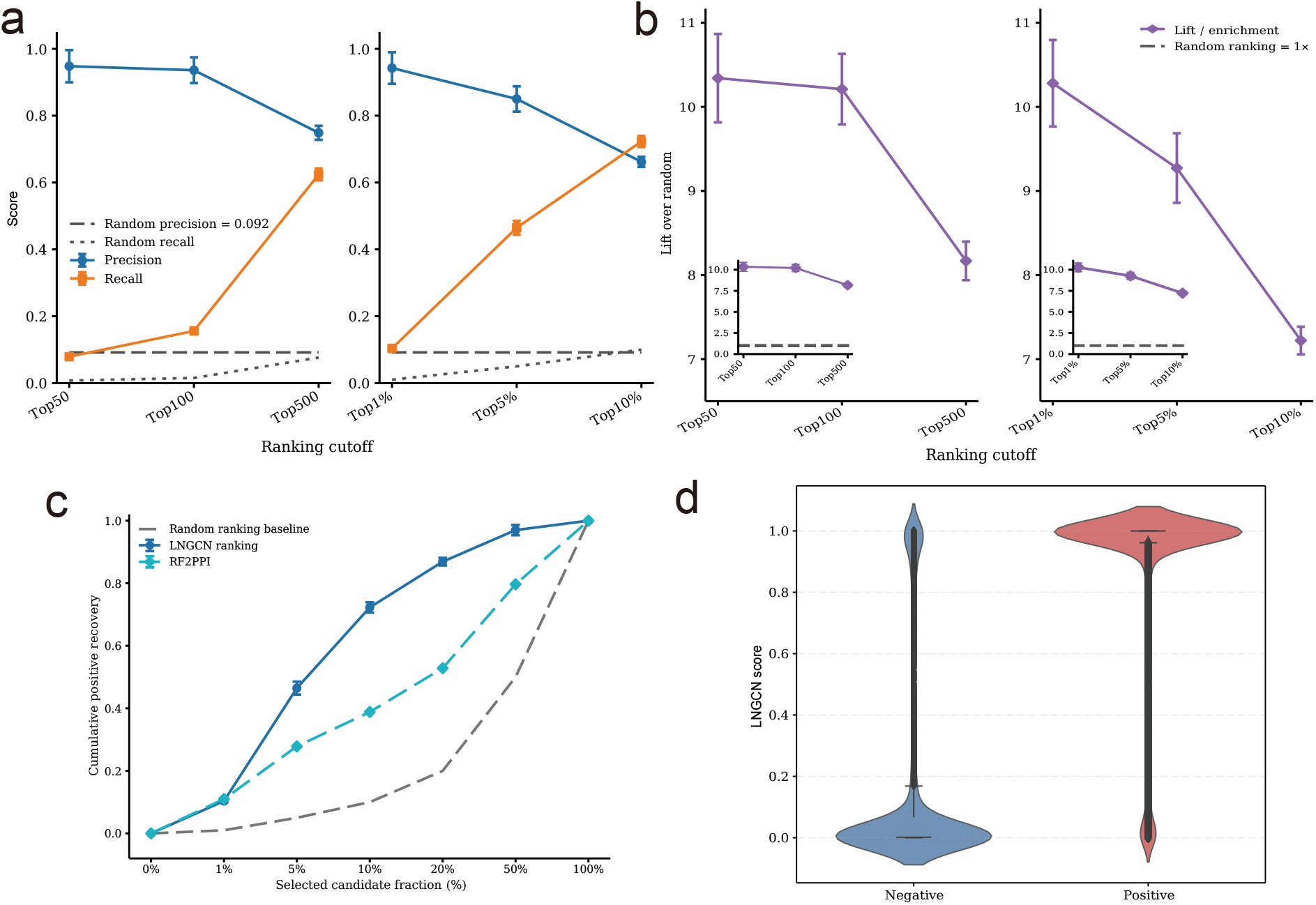
Evaluation of LNGCN candidate ranking under imbalanced PPI and yeast dataset. (A) Precision and recall at fixed candidate numbers (Top 50, Top 100, and Top 500) and screening proportions (Top 1%, *Top*5%, and *Top*10%) on the 1:10 imbalanced human PPI dataset (B) Lift and enrichment relative to random screening at the corresponding ranking cutoffs. (C) Cumulative positive recovery as the selected candidate fraction increases. LNGCN is compared with RF2-PPI and a random-ranking baseline. (D) Distribution of positive and negative protein pairs on the yeast dataset.

The cumulative recovery and enrichment-lift curves consistently placed LNGCN above random ranking and RF2-PPI across screening cutoffs (Figure 4c). The results established that LNGCN not only distinguishes interacting from non-interacting pairs but also concentrates true interactions near the front of highly imbalanced candidate lists, supporting its use for experimentally constrained PPI screening.

### Cross-species evaluation supports transferability candidate ranking

To assess cross-species transferability, the five models trained by cross-validation on the balanced human dataset were directly evaluated on the yeast PPI dataset without retraining. LNGCN achieved a mean AUROC of 0.9352 and a mean precision of 0.8511, indicating that its predictive performance was not restricted to the human training distribution.

LNGCN also retained strong candidate-ranking performance. Across the five models, Precision@50 remained above 0.9800, while NDCG@50 reached 0.9359. At the top-100 cutoff, the mean NDCG increased to 0.9605, demonstrating that true interactions were consistently concentrated near the top of the ranked list even in the cross-species setting.

Score-distribution analysis provided complementary evidence. Violin and density plots (Figure 4d)revealed a clear shift between positive and negative yeast pairs, with limited overlap across much of the score range. Together, the classification, ranking, and distribution results showed that LNGCN captured interaction-related patterns that transferred from human to yeast data. This cross-species performance supported its potential use for PPI candidate prioritization beyond the species represented during model training.

### Continuous-time graph dynamics preserve ranking-relevant representation heterogeneity

Because PPI prioritization depended on maintaining score differences among candidate pairs, we examined whether LNGCN preserved residue-level representation heterogeneity during deep graph propagation. LNGCN was compared with GCN^4^, ResGCN^37^, and GCNII^38^ on the human 1:10 imbalanced dataset and the cross-species yeast dataset.

On the human dataset, representations generated by the discrete graph models became progressively more homogeneous with increasing depth. At the final propagation stage, the node variances of GCN, ResGCN, and GCNII decreased to 3.91, 4.96, and 5.16, respectively, while their Dirichlet energies declined to 0.49, 0.79, and 1.30 (Figure 5a-b). LNGCN showed an initial reduction during neighborhood integration but did not undergo continued representation collapse. Its final node variance remained at 42.20, with a Dirichlet energy of 21.34, substantially exceeding the values obtained by the three comparison models.

**Figure 5:**
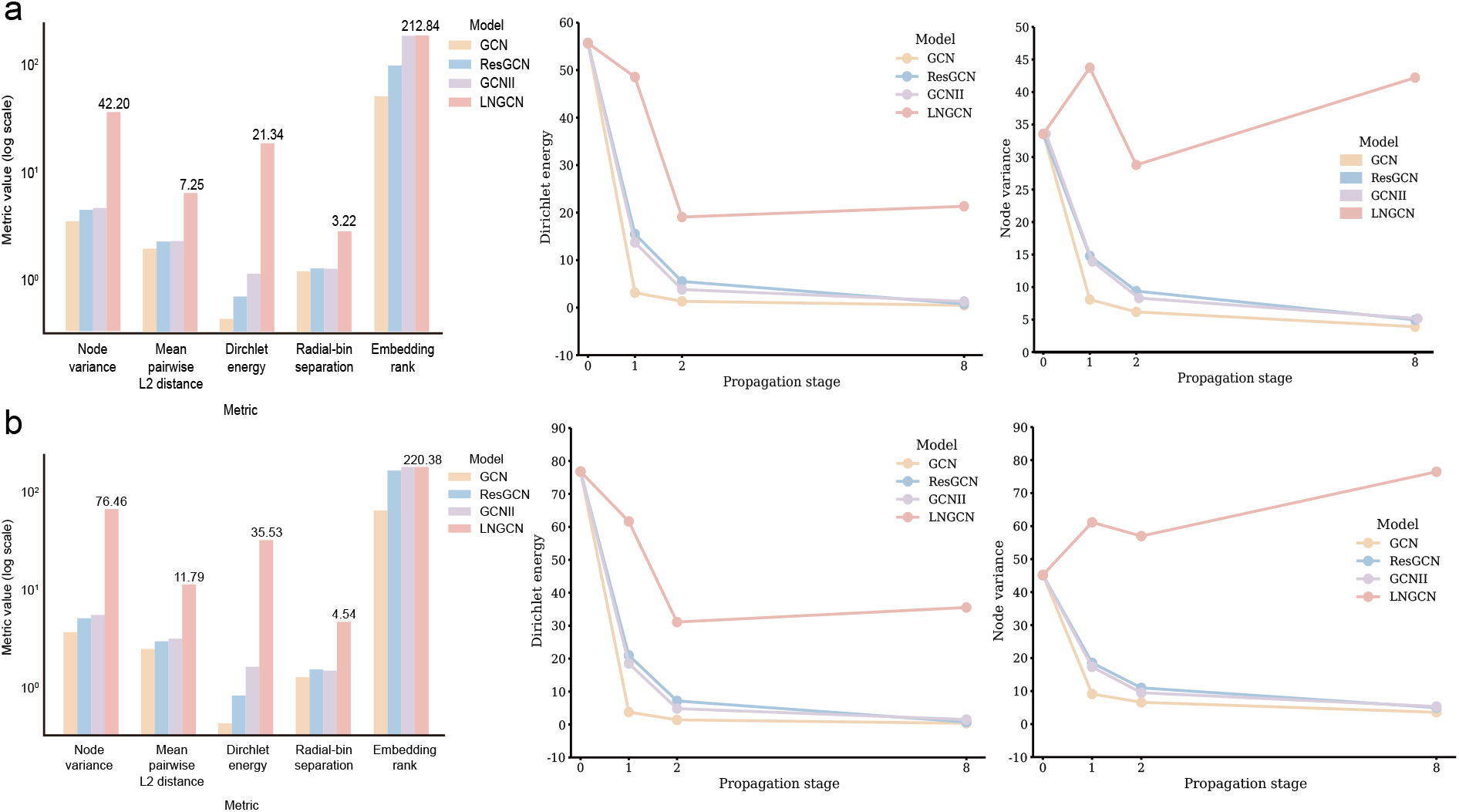
Comparison of representation heterogeneity during graph propagation. (A) Final-stage representation metrics, depth-dependent Dirichlet energy, and node variance for the 1:10 imbalanced human PPI dataset, with residue-level diversity compared to GCN, ResGCN, and GCNII baselines. (B) Corresponding results on the cross-species yeast dataset.

Additional representation metrics yielded a consistent pattern (Figure 5a-b). LNGCN achieved a mean pairwise Euclidean distance of 7.25, a radial-bin separation of 3.22, and an effective embedding rank of 212.84. The larger pairwise distance and radial-bin separation reflected greater differentiation among residue embeddings and between spatially defined residue groups. The high effective rank further indicated that the representation space was not compressed into a small number of dominant dimensions. Although GCNII retained a relatively high effective rank, its node variance, Dirichlet energy, and radial-bin separation remained lower than those of LNGCN, suggesting that global embedding dimensionality alone did not preserve local residue-level differences.

This pattern was reproduced on the yeast dataset. Following continuous-time graph evolution, LNGCN retained a node variance of 76.46 and a Dirichlet energy of 35.53. Its mean pairwise Euclidean distance, radial-bin separation, and effective embedding rank reached 11.79, 4.54, and 220.38, respectively (Figure 5a-b). In contrast, GCN, ResGCN, and GCNII again exhibited rapid decreases in node variance and Dirichlet energy as propagation depth increased.

Across both datasets, LNGCN therefore maintained greater residue-level differentiation than the discrete graph baselines. This representation behavior was consistent with its strong Top-*k* enrichment, NDCG, and cumulative recovery performance on the imbalanced screening dataset, providing a stable representation basis for continuous scoring, calibration, and downstream candidate prioritization.

### Ablation analysis reveals component contributions to prioritization performance

We performed ablation experiments to quantify the contributions of five architectural components. Five variants were evaluated by removing: (A) CfC preprocessing, (B) the graph-driven LNN module, (C) the LNN dense layer, (D) the compensation mechanism, and (E) the combined CfC and LNN modules.

On the human benchmark, the full model achieved an MCC of 0.8936 (Figure 6a). Removing CfC or LNN reduced MCC by approximately 6% and 8%, respectively, whereas removing both modules produced a reduction of approximately 12%. Similar performance changes were observed on the yeast dataset (Figure 6b). A quantitative component analysis further supported this interpretation. On the human test set, the estimated contributions to AUROC were approximately 36% for CfC, 54% for the LNN module, 7% for the LNN dense layer, and 3% for the compensation mechanism (Figure 6c), with a comparable distribution on yeast (Figure 6d). LNN contributed most strongly, while CfC remained important for compressing residue features while preserving radial-distance information.

**Figure 6:**
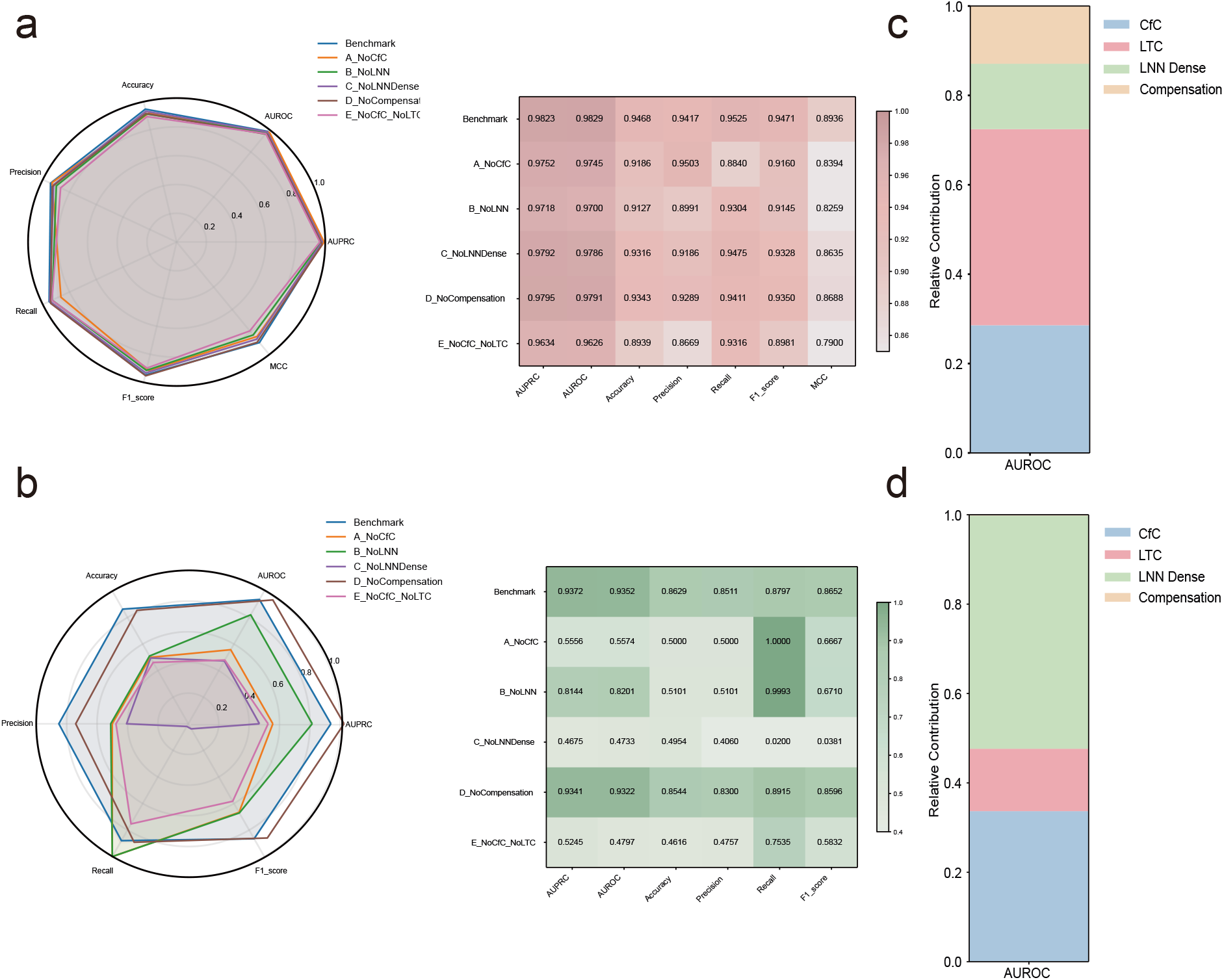
Ablation experiments on benchmark datasets and yeast datasets. (A) Performance radar chart and variance heatmap on the test set in ablation experiments. (B) Performance radar chart and variance heatmap on the yeast dataset in ablation experiments. (C) Quantitative analysis of component contribution on the test dataset. (D) quantitative analysis of component contribution on the yeast dataset.

Ranking ablations on the 1:10 human dataset further revealed losses in candidate enrichment as shown in Figure 7a-b. Removing any individual component reduced Precision@500, Recall@500, and NDCG@500 by more than 10%, while simultaneous removal of CfC and LNN produced reductions exceeding 20%. At the top-10% cutoff, the full model recovered 73.83% of positive pairs with an NDCG of 0.7728. Recall decreased to 63.83% without CfC, 65.33% without LNN, and 61.83% without the LNN dense layer, corresponding to 60, 51, and 72 fewer recovered interactions, respectively. Overall, the ablations establish complementary roles for the continuous-time components in both pair discrimination and early candidate enrichment.

**Figure 7:**
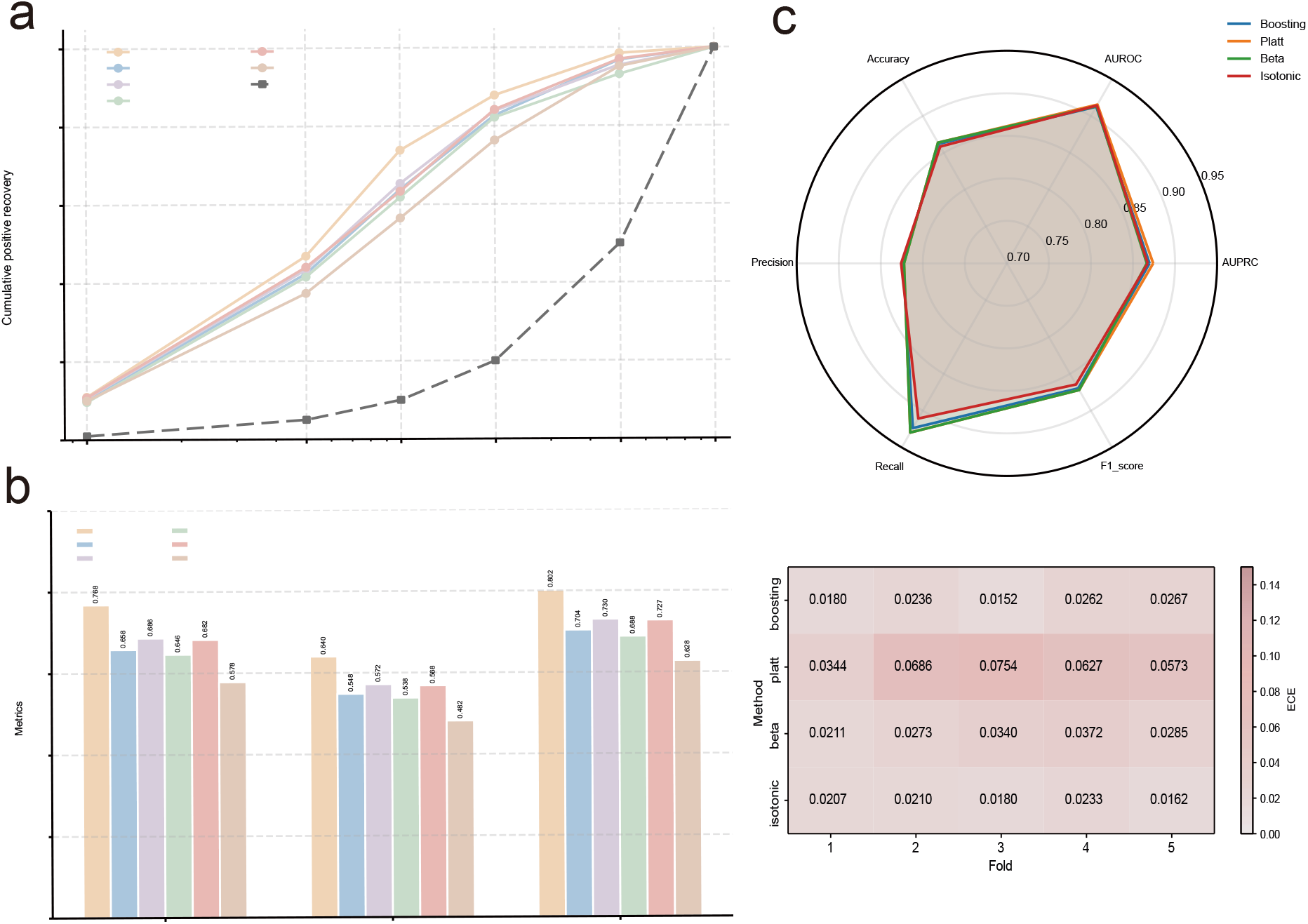
Ablation analysis under imbalanced PPI screening and calibration evaluation. (A) Cumulative positive recovery curves for the complete LNGCN model and its ablation variants across different candidate screening fractions, with random ranking included as a reference baseline. (B) Comparison of Precision@500, Recall@500, and NDCG@500 between the complete LNGCN model and its ablation variants on the 1:10 imbalanced human PPI dataset. (C) Performance comparison of four post hoc calibration strategies. The radar plot summarizes the mean classification metrics across five folds, while the heatmap shows the expected calibration error (ECE) for each calibration method in each fold.

### Hierarchical calibration converts raw predictions into prioritization scores

Because LNGCN was designed for candidate prioritization, its raw outputs were calibrated using a balanced set of 3,000 positive and 3,000 negative pairs^32,35^. Four strategies were evaluated by five-fold cross-validation: Platt scaling^39,40^, Beta calibration^41,42^, Isotonic regression^40,43,44^, and Boosting-based calibration^40,45,46^. All methods retained an AUROC above 0.8971 and an AUPRC above 0.8486 (Figure 7c). Isotonic and Boosting calibration achieved lower expected calibration errors, Beta calibration produced the highest recall (0.9257 ± 0.0364), and Platt scaling achieved the highest AUROC (0.9142 ± 0.0113) and AUPRC (0.8770 ± 0.0178). Because no method was optimal across all criteria, LNGCN used a hierarchical ensemble calibration strategy.

The calibrated output was treated as an empirical prioritization score rather than a biophysical binding probability or a direct estimate of experimental success. Candidate pairs were ranked independently from highest to lowest score. Three operational tiers were defined: low (0–0.4), moderate (0.4–0.7), and high (0.7–1.0). High-scoring pairs were prioritized for validation, whereas moderate-scoring candidates were considered together with orthogonal biological evidence. This calibrated score scale provided a practical basis for threshold selection, candidate comparison, and experimental resource allocation.

### Biological case studies reveal mechanism-consistent candidate prioritization

Having established the early enrichment of true interactions under class imbalance, we next examined whether calibrated LNGCN scores were consistent with known biological organization and could support experimental candidate selection. Three independent biological systems and a TPR case validation case were analyzed. Selected cases also included residue perturbation analysis to identify sequence regions contributing to score variation. These perturbations were used to assess correspondence with literature-supported binding domains, functional regions, or mutation-sensitive sites rather than to predict exact physical contacts.

#### Ternary-complex prioritization in 5W21

The FGF23–FGFR1c–*α*-Klotho ternary complex (PDB: 5W21) was used to evaluate whether LNGCN prioritization scores could reflect known mechanistic relationships within a multiprotein assembly.. In this signaling assembly, *α*-Klotho functions as a non-enzymatic scaffold that facilitates formation of the active FGF23–FGFR1c complex^19^. LNGCN assigned a low score of 0.1613 to FGFR1c–FGF23, whereas FGFR1c–*α*-Klotho and *α*-Klotho–FGF23 received scores of 0.8370 and 0.8017, respectively (Figure 8a). When FGF23 was evaluated against the preassembled *α*-Klotho–FGFR1c complex, its score increased to 0.8238, consistent with the scaffold-dependent organization of the ternary complex.

**Figure 8:**
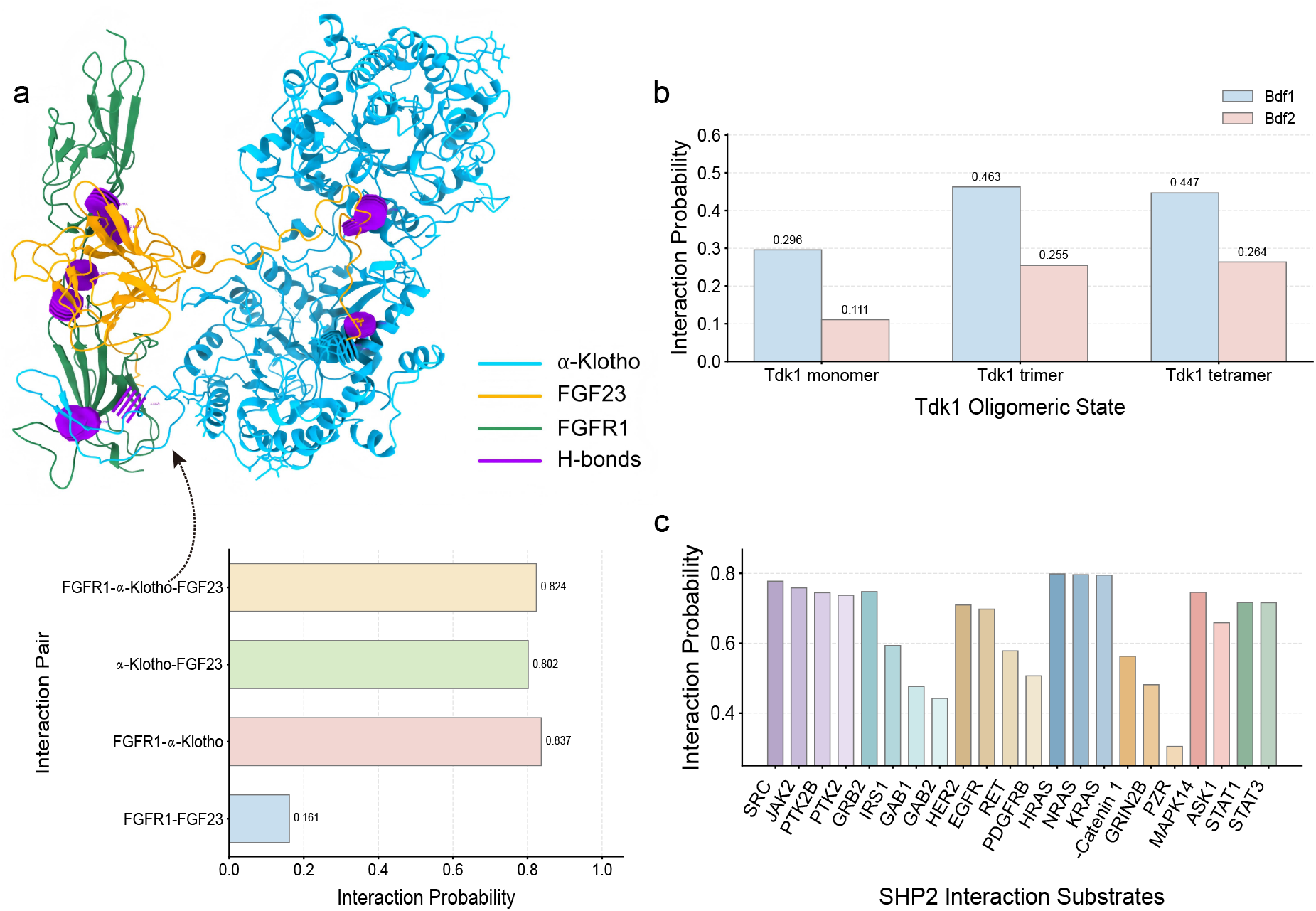
Case studies on PPIs prediction (A) predicted interactions among FGFR1, *α*-Klotho, and FGF23 (B) predicted interaction probabilities between Tdk1 oligomer and Bdf1/Bdf2 (C) predicted interaction probabilities between SHP2 and 22 substrates.

Residue perturbation analysis was performed for the interaction between the preassembled *α*-Klotho–FGFR1c complex and FGF23. High-impact positions included D426 on *α*-Klotho, while influential FGFR1c residues clustered in the D3 domain and near regions associated with heparan sulfate (HS) binding and receptor dimerization. Prominent FGF23 sites were also located near reported HS-binding regions. These patterns connected score variation with structural regions involved in scaffold assembly and HS-mediated dimerization. They do not represent precise contact-site predictions, but suggest that LNGCN prioritization partly depends on structurally plausible regions associated with ternary-complex formation.

#### Oligomeric-state-aware prioritization of Tdk1–Bdf1/Bdf2 interactions

The yeast Tdk1–Bdf1/Bdf2 system was used to examine interaction rankings across oligomeric states and homologous partners. When Tdk1 was evaluated as a monomer, the calibrated prioritization score for the Bdf1-Tdk1 pair reached only 0.2957. This score rose to 0.4627 upon Tdk1 as a trimer, and shifted to 0.4467 in the tetrameric state (Figure 8b). Additional details are provided in Supplemental information S2. This is consistent with reported state-dependent binding behavior^71^. Within the same structural state, Bdf1-associated pairs generally ranked above the corresponding Bdf2 pairs^20^.

The Tdk1 trimer model comprised three Tdk1C fragments connected by artificial linkers. All top-ranked perturbation sites mapped to Tdk1C rather than the linker regions. Recurrent single-residue signals occurred at positions 266–273, 322, and 339, near the distal stalk helix and within the CTD/DUF1773 domain. Pairwise perturbations additionally highlighted Tdk1 residues 245 and 247 within the reported Bdf1-interacting segment 233–255. On Bdf1, residue 408 repeatedly appeared among influential pairs. Although outside the cryo-EM-resolved binding helix at residues 525–547, it lies within the broader Tdk1-binding region 372–554 defined by truncation experiments. These positions were interpreted as perturbation-sensitive, interaction-associated regions rather than exact contact predictions.

#### Phosphorylation-associated prioritization of SHP2 interactions

We next evaluated literature-supported SHP2 signaling interactions involving phosphotyrosine-containing substrates. SHP2 recognizes phosphotyrosine-containing substrates, rendering this system suitable for verifying whether calibrated LNGCN scores align with existing knowledge of PTM-associated signaling pathways. Because ESM-IF1^47^ cannot directly represent phosphorylated structures, unmodified backbone features were supplemented with modified-residue encodings.

Among 22 candidate SHP2-associated pairs, 21 received scores above 0.4 and were supported by published evidence (Figure 8c). High-priority predictions include KRAS, HRAS and NRAS (0.7949-0.7984)^48^, as well as EGFR and HER2 (0.6977 and 0.7096)^49–51^. Strong predictions were also obtained for Src, JAK2, PTK2B, and PTK2^54–58^, alongside GRB2, STAT1/STAT3 and IRS1^59–71^. Full candidate comparisons are provided in Supplemental information S3.

#### TPR candidate prioritization and Co-IP validation

We further evaluate LNGCN’s candidate prioritization performance using nucleoporin TPR as the central target^72^. Known human TPR-interacting proteins were collected from BioGRID^33^, IntAct^34^, and STRING^29^. Protein pairs already present in the model-development datasets were excluded, and no TPR-related data were used for model fine-tuning. Candidate protein pairs were generated by randomly pairing TPR with proteins absent from the above databases and lacking direct literature evidence for TPR binding. Ten additional candidates originating from mass spectrometry interactome data were also incorporated^74^. After filtering for structural availability, 263 known binding partners and 1,340 previously unreported candidates were ranked collectively, forming a total set of 1,603 TPR-centered protein pairs.

Known TPR partners were strongly enriched near the top of the ranked list (Figure 9a-c). A total of 162 out of the 263 known partners (61.6%) exceed the predefined moderate priority score threshold of 0.4, whereas only 84 of the 1,340 unreported candidates (6.3%) exceeded this cutoff. All 39 protein pairs with scores of at least 0.7 were known interactions. The top 10% of the ranking recovered 153 known partners, corresponding to a recall of 58.2%, an apparent precision of 95.0%, and a 5.79-fold enrichment over random selection.

**Figure 9:**
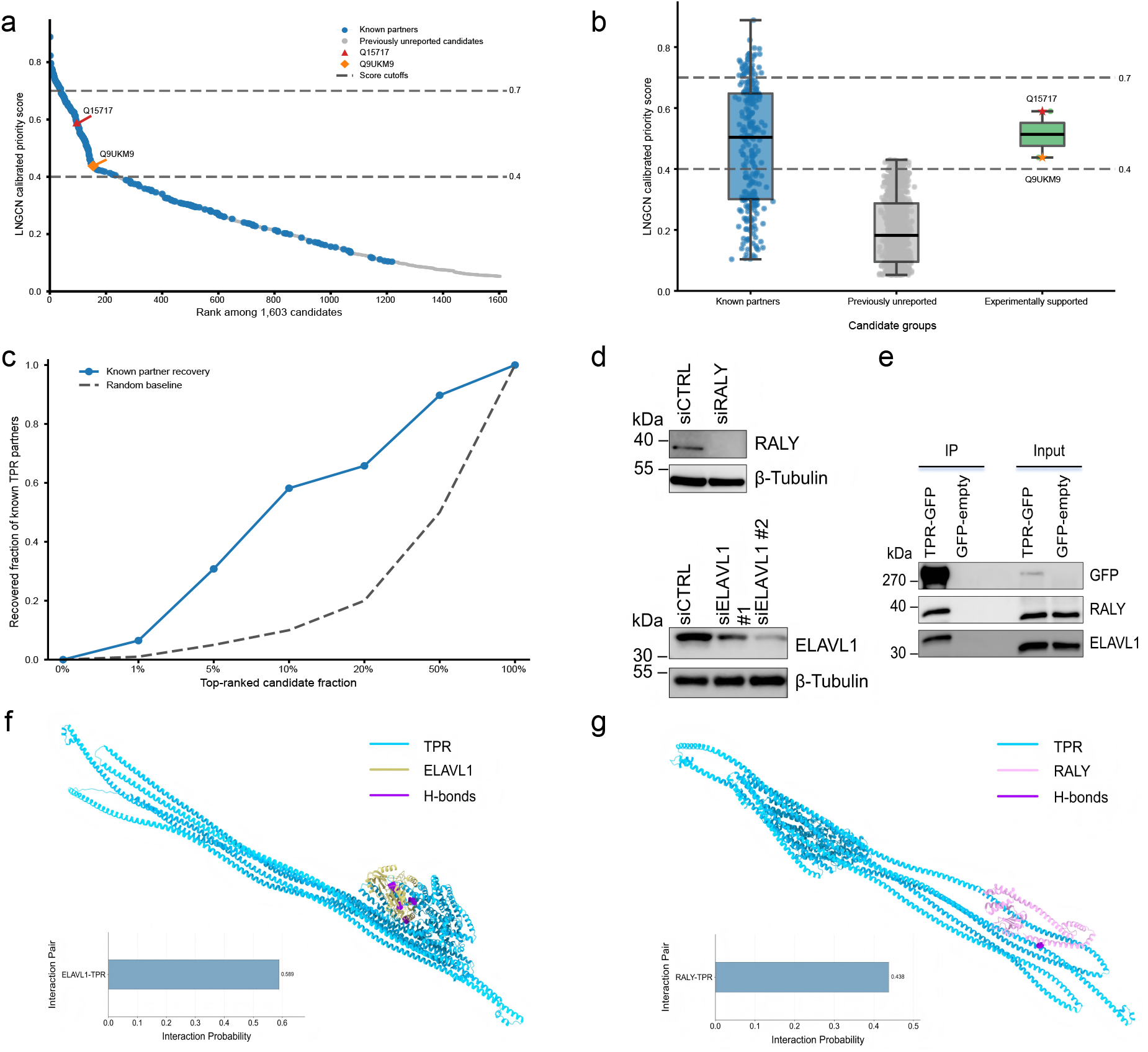
Analysis and validation of TPR-associated PPIs. (A) Calibrated LNGCN priority scores for 1,603 TPR-associated candidate proteins ranked in descending order. (B) Distribution of calibrated priority scores among established TPR partners, candidates without previous direct experimental support, and candidates subsequently supported by experiment. (C) Cumulative recovery of known TPR partners as the proportion of top-ranked candidates increases, compared with a random-ranking baseline. (D) Representative immunoblot analysis of HeLa cell lysates transfected with control siRNA (siCTRL) or siRNA targeting RALY (siRALY) (top), or control siRNA (siCTRL) or two independent siRNAs targeting ELAVL1 (siELAVL1 #1 and siELAVL1 #2) (bottom). Blots validate antibody specificity by showing reduced signal at the expected molecular weight following target knockdown.*β*-Tubulin was used as a loading control. Molecular weight markers (kDa) are indicated on the left. Data are representative of two independent experiments. (E) HeLa cells expressing TPR–GFP or GFP-empty were subjected to anti-GFP immunoprecipitation (IP). Immunoblots of IP and input fractions were probed with antibodies against GFP, RALY, and ELAVL1. RALY and ELAVL1 are enriched in TPR–GFP immunoprecipitates compared with the GFP-empty control. Molecular weight markers (kDa) are indicated on the left. Data are representative of three independent experiments. (F) Schematic diagram of the ELAVL1-TPR interaction. (G) Schematic diagram of the RALY-TPR interactions.

Among the ten mass-spectrometry-derived candidates, the established NUP153–TPR interaction ranked first with a calibrated score of 0.7971^73^. Residue perturbations on TPR were enriched near regions associated with NUP153 or nuclear pore complex binding, including mutation-sensitive positions L458 and M489, while several influential NUP153 residues occurred near reported TPR-binding fragments. For GANP–TPR, this reported interaction received a score of 0.6765. It high-impact TPR positions clustered in N-terminal and nuclear-envelope-binding regions, including the segment surrounding L458.^74^

Within the pool of unreported candidates, ELAVL1–TPR and RALY–TPR receive calibrated scores of 0.5893 and 0.4377, ranking 95th and 154th among all 1,603 pairs. Because neither interaction had previously been supported by direct experimental evidence^29,33,34^, both pairs were selected for follow-up evaluation. We performed knockout-based antibody specificity tests and co-immunoprecipitation assays on both candidate interactors. In HeLa cells transfected with control or target-specific siRNAs, immunoblotting showed that the RALY and ELAVL1 signals at their expected molecular weights were markedly reduced after knockdown (Figure 9d), supporting the specificity of the corresponding antibodies. Anti-GFP co-immunoprecipitation further demonstrated greater enrichment of RALY and ELAVL1 in TPR-GFP precipitates than in the empty-vector GFP control (Figure 9e). Input immunoblots confirmed that both proteins were present in the cell lysates. Structural schematics of these validated interaction pairs are presented in Figure 9f-g. RF2-PPI^32^ assigned lower probabilities to ELAVL1–TPR and RALY–TPR (0.2090 and 0.2861), whereas LNGCN placed both higher in the candidate list before Co-IP validation.

Across the biological cases, LNGCN scores captured known complex organization, oligomeric-state dependence, and signaling relationships, while the TPR application connected candidate ranking with experimental follow-up. Residue perturbation analyses also suggested that LNGCN prioritization scores were partly attributable to biologically plausible residue-level regions, rather than purely driven by model training.

## DISCUSSION

In this study, we developed LNGCN as a candidate prioritization framework for experimental PPI screening. Rather than treating PPI prediction solely as a binary classification task, LNGCN generates stable, continuous, and calibrated scores for large collections of candidate protein pairs to guide selection of the most promising interactions. The framework combines multi-modal sequence and structural features, distance-aware continuous-time residue graph modeling, symmetric protein pair fusion, and raw score calibration. It was evaluated through balanced classification, ranking under class imbalance, cross-species transferability, representation heterogeneity, ablation analysis, and biological case studies.

On the balanced human benchmark, LNGCN achieved mean AUPRC and AUROC values above 0.98 with limited variation across five folds. Performance also remained useful under the 1:10 imbalanced setting, which better resembles practical screening. Approximately 84.45% of candidates within the top 5% were positive. The Top-K, NDCG, enrichment-lift, and cumulative recovery results therefore addressed the need to identify informative candidates from a predominantly negative search space. Direct application of the human-trained models to yeast data also retained strong classification and early-ranking performance, suggesting that LNGCN captured transferable interaction-related patterns rather than only characteristics specific to the human training distribution.

This ranking performance was accompanied by preservation of residue-level representation heterogeneity. Deep graph convolution could progressively homogenize node features, reducing the distinctions needed to separate closely ranked candidate pairs. LNGCN instead retained higher node variance, Dirichlet energy, pairwise embedding distance, radial-bin separation, and effective embedding rank after neighborhood integration on both human and yeast datasets. It provided a plausible representational basis for the differentiated continuous scores. Removing CfC or the graph-driven LNN module reduced classification and ranking performance, while removing both produced the largest decline. CfC-based feature processing and continuous-time graph dynamics therefore appear to make complementary contributions to candidate scoring.

Calibration further improved the operational value of the outputs. Because no individual method was optimal across discrimination, recall, and calibration error, LNGCN used a hierarchical ensemble calibration. The resulting values should be interpreted as empirical prioritization scores, not binding affinities or biophysical probabilities. The LNGCN backbone determined the relative ordering of candidate pairs, whereas calibration makes the score range more suitable for comparison, and allocation of validation resources. The objective was not to estimate molecular affinity, but to convert model outputs into a usable screening scale.

The biological case studies examined whether the calibrated rankings remained consistent with known organizational principles. In the 5W21 system, the scores reflected scaffold-dependent assembly of the FGF23–FGFR1c–*α*-Klotho complex. In the Tdk1 system, rankings varied with oligomeric state and distinguished Bdf1 from Bdf2. The SHP2 analysis placed literature-supported signaling partners among the higher-scoring candidates. The strongest application-level evidence came from the TPR study. Across 1,603 jointly ranked TPR-centered pairs, established partners were strongly enriched near the top, with the top 10% recovering 58.2% of documented interactions. ELAVL1–TPR and RALY–TPR were both prioritized above the predefined experimental threshold and subsequently received the first pair-specific support from co-immunoprecipitation. The analysis of partial residue perturbations further localized score sensitivity to literature-supported domains, functional regions, and mutation-sensitive segments. These results should not be interpreted as evidence of direct binding or precise contact-site prediction, but they reduce the concern that prioritization reflects only memorized training-set associations.

Several limitations remain. LNGCN did not explicitly model condition-dependent factors such as subcellular localization, cellular state, or disease context. Cross-species evaluation was limited to yeast, and broader testing across additional organisms will be needed. The current framework also relied on static structures and therefore cannot fully represent conformational transitions, transient interactions, or interactions mediated by intrinsically disordered regions. Future work could incorporate conformational ensembles or molecular-dynamics trajectories as time-indexed residue graphs, allowing continuous dynamics to model structural evolution and interaction propensity. Finally, perturbation-derived residues remain model-dependent hypotheses requiring validation through functional experiments. And these limitations may provide clear directions for future development of the framework.

Overall, LNGCN connected residue-level structural representation, continuous-time graph dynamics, calibrated scoring, and experimental candidate selection within a single framework. Its benchmark performance, early enrichment under class imbalance, cross-species transferability, preservation of representation heterogeneity, together with biologically relevant cases and experimentally validated candidate identification, demonstrate its potential as a scalable tool for translating PPI predictions into practical prioritization screening.

## METHODS

### Residue-level graph construction

To precisely characterize the three-dimensional (3D) structure of proteins, we downloaded the full-length structural data of target proteins predicted by AlphaFold^30^ from the UniProt database, and extracted their 3D atomic coordinate information, i.e., the precise spatial position information of the *Cα* atom of each residue. We adopt a residue contact map as the representation of protein 3D structure, with the graph formally defined as *G* = (*V, E*), where V represents all residue nodes in the protein, and *E* represents the contact edges between residues. For any two residues *i* and *j*, if the Euclidean distance between their *Cα* atoms is less than a threshold *d*_*c*_ (*d*_*c*_ = 8 Å), they are considered to be in contact, which is mathematically expressed as:

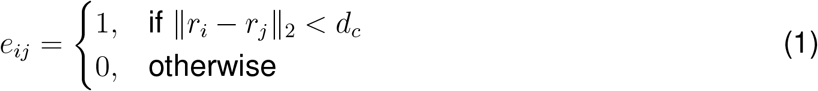

where, *r*_*i*_ and *r*_*j*_ correspond to the 3D coordinate vectors of the *Cα* atoms in residues *i* and *j*, respectively.

To accomplish the quantitative construction of the aforementioned residue contact map, we first calculated the distance matrix for all residue pairs in the protein based on atomic coordinates, with the calculation formula as follows:

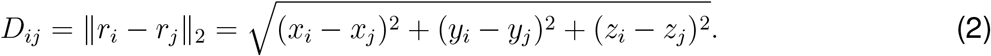

Then, the adjacency matrix A is generated by thresholding:

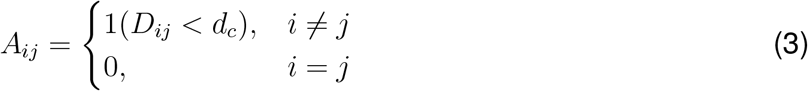

The final constructed graph was an undirected graph. We implemented the construction of this undirected graph using the Deep Graph Library (DGL)^75^, and simultaneously added self-loop connections to each node to enhance the retention and transmission of the node’s own information.

### 1 Node features

For each residue, we constructed a 1803-dimensional multimodal feature vector by integrating ESM-2 sequence embeddings^22^, ESM-IF1 structural embeddings^47^, DSSP secondary-structure descriptors^24^, and relative solvent accessibility features^25–28^.

Protein sequences were encoded using the pretrained ESM-2 model (*esm*2_*t*33_650*M*.*pt*, 650M parameters)^22^. Each residue was represented by a 1280-dimensional embedding, producing an *L* × 1280 feature matrix for a protein of length *L*. Residue-level structural representations were extracted using ESM-IF1^47^. Given the backbone coordinates (*N, Cα, C*), ESM-IF1 generated a 512-dimensional embedding for each residue. Secondary structures were assigned using DSSP^24^. Its nine structural states were encoded as 9-dimensional one-hot vectors. These explicited conformational descriptors complemented the learned ESM-IF1 representations. Residue solvent exposure was calculated using FreeSASA^25^. The absolute solvent-accessible surface area was normalized by the theoretical maximum value of each amino-acid type obtained from AAindex^26–28^, yielding a relative solvent accessibility (RSA) value between 0 and 1. Based on these RSA values, we further developed a binary “interaction propensity” feature: a residue was labeled as 1 when its RSA exceeded 0.25, and 0 otherwise, so as to explicitly mark highly exposed residues that were potentially located at the PPI interface.

To explicitly encode global spatial position, we additionally introduced a normalized residue radial distance feature. Let *e*_*i*_ denote the *Cα* coordinate of residue *i*, and let 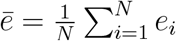 denote the protein center. The residue-to-center distance and its normalized value were defined as:

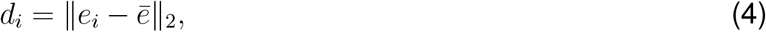

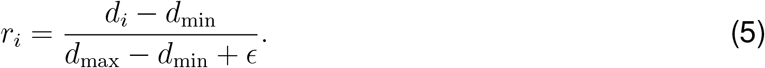

Here, *ϵ* = 10^−8^ ensures numerical stability, and *r*_*i*_ ∈ [0, 1] distinguishes relatively central from peripheral residues. Unlike the 1803-dimensional static feature vector, *r*_*i*_ was retained separately as an explicit spatial driving signal for the distance-aware dynamic module.

Especially, RSA and residue radial distance feature provide complementary spatial information. RSA reflects local solvent exposure determined by the surrounding molecular surface, whereas residue radial distance feature describes global position relative to the protein center. They are not interchangeable. Residues at similar radial positions may differ in exposure because of grooves, cavities, or irregular protein geometry, while residues with similar RSA values may occupy distinct global regions. Their joint use therefore allows LNGCN to distinguish local accessibility from global spatial organization during interaction-oriented candidate scoring.

#### CfC-based feature preprocessing

The initial node feature vector contains high-dimensional heterogeneous information. To reduce redundancy while retaining structurally useful signals, LNGCN uses a closed-form continuous-time network (CfC) as a gated preprocessing module. CfC provides an efficient time-dependent gating mechanism^18^. Its generic form interpolates between a dynamic branch and a static branch:

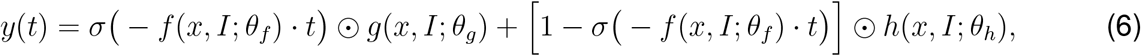

where *σ* is Sigmoid function.

In LNGCN, CfC is adapted as a feedforward gated projection for residue graphs. The input node vector *x*_*i*_ ∈ ℝ^1804^ is divided into a 1803-dimensional base feature and a one-dimensional radial distance *r*_*i*_. The base features pass through three CfC layers with pseudo-time *τ* ^(*l*)^ = *l/L*. For layer *l*, the latent representation 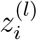 is projected into gating coefficients and dynamic/static candidate states:

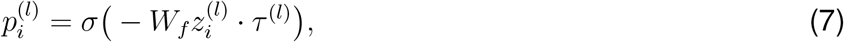

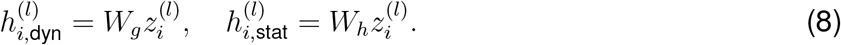

The output is obtained by element-wise interpolation:

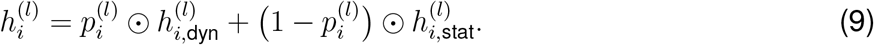

The final CfC output compresses the base representation to 256 dimensions and appends the unchanged radial distance:

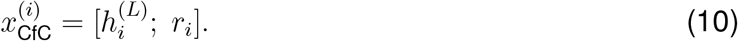

This step reduces feature noise while preserving the spatial variable used for distance-aware graph evolution.

#### Distance-aware graph-driven LNN module

We extend the continuous-time formulation to each residue node in the protein graph:

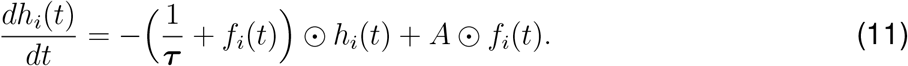

where each residue has a node-specific driving term *f*_*i*_ = *f* (*h*_*i*_, *N* (*i*), *r*_*i*_; *θ*). Here, *N*(*i*) denotes the graph neighborhood of node *i, r*_*i*_ is its normalized radial distance, and *θ* contains the learnable parameters.

The normalized radial distance *r*_*i*_ ∈ [0, 1] is first projected into the node-feature space:

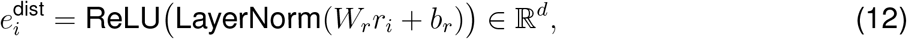

At ODE step *k*, let *H*^(*k*)^ ∈ ℝ^*N* ×*d*^ denotes the current residue representations. Graph-neighborhood information is aggregated as

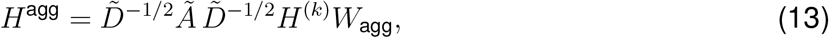

where *Ã*= *A*_adj_ + *I* includes self-loops and 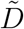 is its degree matrix. The *i*th row of *H*^agg^ is denoted by 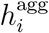.

For each node, the aggregated graph feature and radial-distance embedding are concatenated and projected back to *d* dimensions:

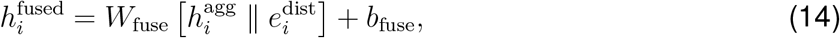

where *W*_fuse_ ∈ ℝ^*d×2d*^. A channel-wise gate is then applied to generate the node-specific driving term:

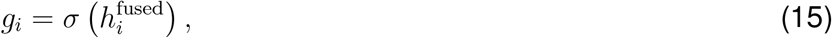

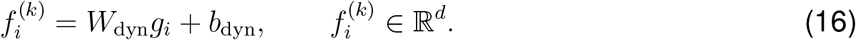

This formulation allows graph-neighborhood information and residue-specific spatial position to jointly determine the driving term 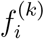, producing spatially heterogeneous node dynamics.

The continuous dynamics are discretized using implicit Euler:

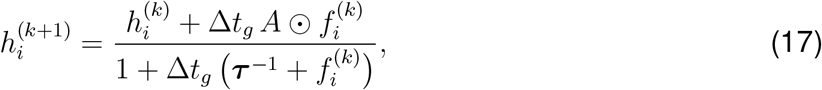

The parameter *A* is initialized as ones and constrained within (0, 10), while ***τ*** ∼ *U* (0.5, 5). This implicit scheme is numerically stable^76^. The solution trajectory is continuously differentiable with respect to the radial distance under standard ODE sensitivity analysis^77^, supporting gradient-based learning of spatially dependent interaction features. Formal stability analysis and propositions are provided in Supplemental information S4.

After *S* rounds of ODE unfolding, residue states are stacked as 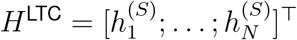. Multi-hop graph convolution and residual fusion are then applied to capture local and extended structural neighborhoods. This module took *H*^LTC^ as input and extracted structural features through three branches at different scales.

Treat the ODE output as a zero-order characteristic and denote it as

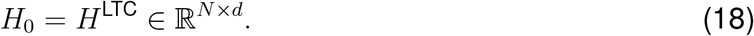

Then, in the first-order branch, a graph convolution layer followed by ReLU activation was applied to *H*_0_, which captured 1-hop neighborhood information, written as:

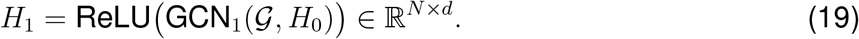

Furthermore, on the second-order branch, we applied graph convolution again with *H*_1_ as input, followed by ReLU activation, to obtain feature *H*_2_ containing 2-hop neighborhood information, written as:

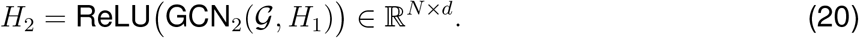

Subsequently, the features from the three scales were concatenated along the channel dimension and fused back to the original dimension through a linear layer:

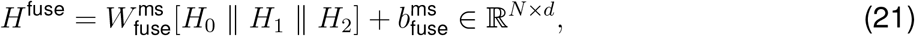

where 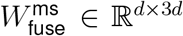 is the multi-scale fusion weight matrix. To enhance gradient propagation and training stability, we introduced an auxiliary graph convolution layer GCN_aux_ (an independent single-layer GCN that learns complementary structural features from *H*_0_, providing a parallel feature extraction path alongside the multi-scale fusion), residual connections and layer normalization to integrate the multi-scale fusion with auxiliary features, obtaining the final node-level output:

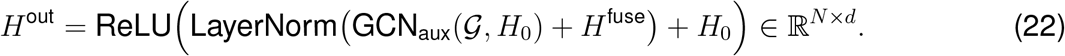

Based on the node-level representations *H*^out^ obtained above, we derived the graph-level representation of the protein by performing average pooling over all node features:

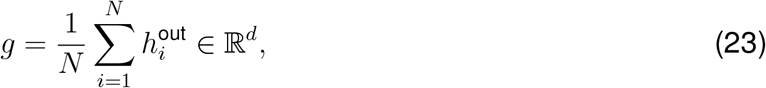

where 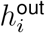 denotes the *i*^*th*^ row of *H*^out^, and *g* corresponds to the fixed-dimensional graph-level embedding vector for the entire protein.

#### Protein-pair fusion and LNN dense classifier

For a candidate protein pair, the graph-level representations **g**_1_ and **g**_2_ are combined using a symmetric fusion strategy:

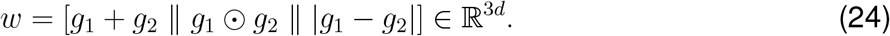

This representation captures shared patterns, multiplicative compatibility and absolute differences while remaining invariant to protein order.

The fused vector is passed into two LNN dense layers rather than conventional fully connected layers. Starting from *w*^(0)^ = *w*, the hidden state evolves for *N* = 10 discretization steps with *δt* = 0.1:

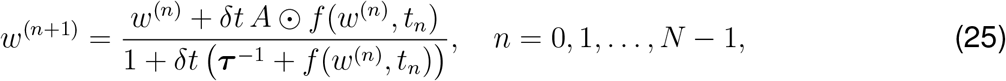

The driving function concatenates the current state and pseudo-time:

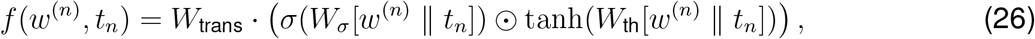

A final linear projection generates binary classification logits:

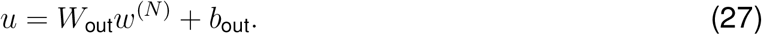

#### Probability calibration

For experimental screening, calibrated probabilities are more useful than raw classification scores because they provide more interpretable confidence estimates for candidate ranking and threshold-based selection. LNGCN therefore incorporates a post-processing calibration module. We evaluated four calibration methods, including Platt calibration^39,40^, Beta calibration^41,42^, Isotonic calibration^40,43,44^, and Boosting calibration^40,45,46^, with detailed procedures provided in Supplemental information S5.

Let *s*_*i,k*_ denote the raw model score assigned to candidate protein pair *i* by the LNGCN model trained in fold *k*, where *k* = 1, … , *K* and *K* = 5. Let *g*_*k,c*_() denote calibrator *c* fitted in fold *k*, where *C* = {Platt, Beta, Isotonic, Boosting} is the fixed set of calibration methods used in each fold. The calibrated probability generated by calibrator *c* for candidate pair *i* in fold *k* was defined as

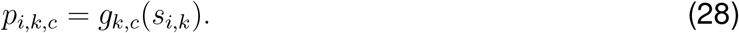

To reduce dependence on the assumptions of any single calibration method, we adopted a hierarchical ensemble strategy across calibration methods and cross-validation folds. First, predictions from the four calibrators within each fold were averaged:

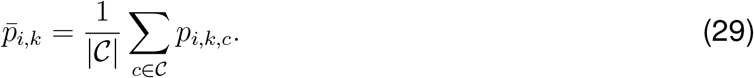

Second, cross-fold bagging was applied to obtain the final calibrated interaction probability:

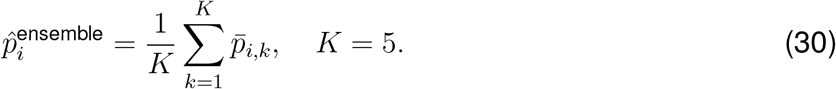

Cross-fold averaging was used to reduce sensitivity to individual training splits and provide more stable predictions for external candidates. The final output 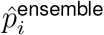 was used as the calibrated interaction probability for candidate ranking.

#### Experimental Methods

##### Cell culture

HeLa cells (European Collection of Authenticated Cell Cultures) were maintained in Dulbecco’s Modified Eagle’s Medium (DMEM) (Sigma-Aldrich, #D5671) supplemented with 10% FBS, 1 mM sodium pyruvate, 1× MEM non-essential amino acids, 2 mM GlutaMAX, and 100 U/mL and 100 *µ*g/mL penicillin-streptomycin at 37°C with 5% CO_2_. All cell lines were routinely tested and confirmed mycoplasma-negative. Complete medium composition is in Supplemental information S6.

##### Transfections

HeLa cells (six million/dish) were seeded on eight 145-mm dishes (four dishes per condition) and transfected the following day with pcDNA4/TO-TPR-2siR-EGFP (for TPR expression) or empty pcDNA4/TO-EGFP (negative control) using Neofect DNA Transfection Reagent. Alternatively, siRNA transfections were performed using Lipomaster 3000 (Vazyme, #TL301-02) at a 25 nM final concentration. Detailed protocols are in Supplemental information S6.

##### Immunoprecipitation

Immunoprecipitation followed a published protocol (Collis et al., 2007)^79^. Transfected cells were washed with cold 1x PBS, harvested, and centrifuged (300 x g, 4°C, 5 min). Cell pellets were resuspended in ice-cold lysis buffer and incubated on ice for 30 min, pipetting to reduce viscosity every 10 min. Lysates were clarified by centrifugation (16,000 x g, 4°C, 10 min). Following protein quantification via a BCA assay (Yeasen, #20201ES90), equal protein amounts were incubated overnight at 4°C with anti-GFP magnetic beads (MCE, HY-K0246). The next day, beads were magnetically separated, washed three times with dilution buffer, and proteins were eluted in 1× SDS sample buffer at 95°C for 10 min. The lysis, dilution and SDS sample buffer compositions are detailed in Supplemental information S6.

##### Immunoblotting

Whole-cell extracts were prepared in 1× SDS sample buffer, sonicated (Xiaomei Chaosheng Instruments, XM-650T, China), and centrifuged (16,000 × g, 4°C, 30 min). Protein concentrations were determined using a BCA assay. Proteins were separated via SDS-PAGE and transferred to PVDF membranes (Millipore, #IPFL00010). Membranes were blocked in 5% milk-PBST for 30 min at room temperature. Primary antibodies were diluted in 1% (w/v) milk-PBST and incubated overnight at 4 °C.Secondary antibodies were diluted in the same buffer and incubated for 1 h at room temperature. Signals were detected using FG Super Sensitive ECL Luminescence Reagent (Meilunbio, #MA0186). Details of antibodies and SDS sample buffer are available in Supplemental information S6.

## RESOURCE AVAILABILITY

### Lead contact

Requests for further information and resources should be directed to and will be fulfilled by the lead contact, Ying Chi

### Materials availability

This study did not generate new unique reagents.

### Data and code availability

The source datasets used in this study are publicly available from the STRING database (https://string-db.org/), UniProt database (https://www.uniprot.org/), Negatome database (https://mips.helmholtz-muenchen.de/proj/ppi/negatome/), BioGRID database (https://thebiogrid.org/), and AlphaFold Database (https://alphafold.ebi.ac.uk/). The processed datasets generated in this study have been deposited in github (https://github.com/yueming521/LNGCN_main.git).

The custom code for the LNGCN model, data processing, and analysis pipelines used in this study are publicly available on GitHub (https://github.com/yueming521/LNGCN_main.git) to ensure long-term availability.

## ACKNOWLEDGMENTS

This work was supported by the International Scientific Research Cooperation Seed Fund of International Campus, Zhejiang University, for the project entitled “Artificial Intelligence-Assisted Molecular Dynamics Simulation and Its Application and Validation in BK Channels”. The authors thank all members of the lab for their support.

We would like to thank students Make Zhao (Zhejiang University), Jingjing Zhao (Zhejiang University), Yifei Yao (Zhejiang University), Xinxin Zhang (Zhejiang University), Qiufan Xu (Zhejiang University) and Su Shen (Zhejiang University) for their valuable technical support during the data analysis. We also acknowledge the support from the Cross Lung Research Alliance (CLRA) Hub (where Y.K, Y.C, Y.X, J.P, and Y.Q are affiliated), the AI for Life-Course Epidemiology of Stress, Infection, and Healthy Aging (SHENG AI) Hub (where Y.C, Q.Q, and R.F are affiliated), and the Center for Biomedical System & Informatics, where all authors of this study other than Y.K, M.K, and Y.H are affiliated.

## AUTHOR CONTRIBUTIONS

Y.X. , Y.C. , Y.K. and M.K. conceived the project. Y.C. , Y.K. and M.K. supervised the entire study. Y.C. and Y.K. supervised all model-related work, and M.K. supervised all biological experiments. Y.X. co-designed the model architecture with J.P. , J.L. and Y.X., J.P. and Y.Q. jointly designed the data features and carried out model construction and ablation experiments. Y.X., Y.Z., Y.H. and J.X. explored specific application scenarios and conducted downstream functional analyses. Y.Z. completed the experimental validation of PPIs. R.F. and Q.Q. generated all result visualisations and provided additional support for code implementation and debugging. Y.X. drafted the model section, whilst Y.Z. drafted the experimental section. All authors contributed comments and reviewed the final version.

## DECLARATION OF INTERESTS

The authors declare no competing interests.

## SUPPLEMENTAL INFORMATION INDEX

### Document S1

Supplemental Tables S1–S4, extended biological analyses, theoretical propositions and proofs, probability calibration details, experimental procedures, and supplemental references

## Notes

### Competing Interest Statement

The authors have declared no competing interest.

### Summary of Updates

This version of the manuscript has been substantially revised to strengthen its positioning as a protein protein interaction candidate prioritization framework. Ranking based evaluations were added for both the imbalanced human dataset and the cross species generalization dataset, including early enrichment and candidate recovery analyses. Additional analyses and a new figure were included to evaluate whether residue level representations retained sufficient heterogeneity during graph propagation, providing empirical evidence that the proposed continuous time dynamics reduce representation homogenization. Ranking focused ablation experiments were also added to quantify the contributions of the major model components under class imbalanced conditions. To improve model interpretation, residue perturbation analyses were conducted for selected biological cases to assess how individual residues and residue pairs influence interaction priority scores. In addition, the TPR centered application was expanded into a complete case study, including candidate selection, full ranking results, recovery of known interaction partners, prioritization of previously unvalidated candidates, and experimental validation. The corresponding Methods, Results, figure legends, supplemental information, and Discussion sections were revised to reflect these additions and to clarify the distinction between interaction prioritization scores and physical binding affinity.

