## supplemental information for "LNGCN: a distance-aware continuous-time graph framework for PPI prioritization in experimental screening"

### S1 Evaluation of the overall performance of the LNGCN model on the benchmark dataset, imbalanced datasets, and cross-species test

Table S1: The model's performance on the benchmark dataset using 5-fold cross-validation. The metrics include AUPRC, AUROC, Accuracy, Precision, Recall, F1-Score, and MCC. Each metric is reported for each fold of the cross-validation.

| Fold | AUPRC | AUROC | Accuracy | Precision | Recall | F1-Score | MCC |
|---|---|---|---|---|---|---|---|
| 1 | 0.9826 | 0.9826 | 0.9443 | 0.9417 | 0.9471 | 0.9444 | 0.8886 |
| 2 | 0.9724 | 0.9713 | 0.9168 | 0.9291 | 0.9024 | 0.9156 | 0.8339 |
| 3 | 0.9829 | 0.9827 | 0.9441 | 0.9386 | 0.9503 | 0.9444 | 0.8883 |
| 4 | 0.9827 | 0.9830 | 0.9445 | 0.9362 | 0.9540 | 0.9450 | 0.8892 |
| 5 | 0.9823 | 0.9829 | 0.9468 | 0.9417 | 0.9525 | 0.9471 | 0.8936 |

Table S2: Performance on imbalanced datasets using 5-fold cross-validation

| Fold | AUPRC | AUROC | Accuracy | Precision | Recall | F1-Score | MCC |
|---|---|---|---|---|---|---|---|
| 1 | 0.7699 | 0.9481 | 0.9502 | 0.7299 | 0.7250 | 0.7274 | 0.7000 |
| 2 | 0.7343 | 0.9423 | 0.9346 | 0.6129 | 0.7783 | 0.6858 | 0.6556 |
| 3 | 0.7556 | 0.9425 | 0.9291 | 0.5827 | 0.7983 | 0.6737 | 0.6449 |
| 4 | 0.7396 | 0.9420 | 0.9384 | 0.6455 | 0.7283 | 0.6844 | 0.6519 |
| 5 | 0.7619 | 0.9385 | 0.9429 | 0.6692 | 0.7450 | 0.7050 | 0.6747 |

Table S3: Performance on the cross-species test using 5-fold cross-validation

| Fold | AUPRC | AUROC | Accuracy | Precision | Recall | F1-Score | MCC |
|---|---|---|---|---|---|---|---|
| 1 | 0.9372 | 0.9325 | 0.8602 | 0.8493 | 0.8759 | 0.8624 | 0.7209 |
| 2 | 0.9379 | 0.9334 | 0.8558 | 0.8370 | 0.8836 | 0.8597 | 0.7126 |
| 3 | 0.9373 | 0.9347 | 0.8616 | 0.8450 | 0.8857 | 0.8649 | 0.7241 |
| 4 | 0.9347 | 0.9319 | 0.8581 | 0.8391 | 0.8860 | 0.8619 | 0.7172 |
| 5 | 0.9372 | 0.9352 | 0.8629 | 0.8511 | 0.8797 | 0.8652 | 0.7262 |

### S2 Detailed Mechanistic Analysis of Tdk1 Tetramer Self-Inhibition

The predicted probability of Bdf1 forming a tetramer was 0.4467, showing a slight decrease (approximately 0.02) compared to the trimer (0.4627), although both fall within the moderate interaction range. According to structural studies by Hua et al., the natural Tdk1 tetramer generated steric hindrance effects by forming a 4-helix coiled-coil conformation, thereby inhibiting Bdf1's accessibility to its binding interface[1].

However, the tetramer construction in our study employed flexible linker peptides to concatenate four monomers into a single chain for structure prediction input, indicating that simply concatenating into a single-chain tetramer may not fully recapitulate the special 4-helix coiled-coil in the native structure. Consequently, the predicted probability difference between tetramers and trimers failed to adequately reflect the self-inhibition effect documented in the literature. Nevertheless, the slight downward trend in tetramer probability relative to trimer probability still reflected a tendency toward interaction suppression to some extent.

### S3 Detailed SHP2-Substrate Interaction Analysis by Protein Families

Among small GTPases of the Ras family, KRAS, HRAS, and NRAS obtained the highest predicted probabilities ($0.7949 - 0.7984$). These findings were highly consistent with the study by Bunda et al., which confirmed that SHP2 promoted Ras activation through dephosphorylating its Tyr32 site, and that inhibiting this process effectively suppresses tumorigenesis[2]. These three Ras subtypes, as small-molecule GTPases, serve as pivotal switches in cell proliferation, differentiation, and survival signaling pathways. The predicted high-efficiency dephosphorylation of Ras by SHP2 validated this phosphatase's crucial regulatory role in activating the Ras pathway.

In the tyrosine kinase family, the predicted probabilities for EGFR and HER2 were $0.6977$ and $0.7096$, respectively. Zhou and Agazie elucidated the molecular mechanisms by which SHP2 promoted HER2-induced signal transduction and cellular transformation[3]. Hartman et al. further revealed that specific amino acid sequence environments at EGFR and HER2 phosphorylation sites determined their selective binding to SHP2 active sites[4]; Zhan et al. confirmed that SHP2 was essential for EGFRvIII-mediated malignant transformation in glioblastoma[5]. These findings provided robust support for the phosphorylation-dependent regulatory relationship between EGFR/HER2 and SHP2. The predicted probability for PDGFRB was $0.5068$. Research by Bennett et al. demonstrated that SHP2 couples PDGFRB to the Ras pathway[6], providing evidence for the predicted interaction. The predicted probability for the RET receptor was $0.5782$. Perrinjaquet et al. also confirmed that SHP2 contributes to GDNF neurotrophic activity through direct binding to phospho-Tyr687 in the RET receptor[7].

In the scaffold protein family, the predicted probability for Src was $0.7776$, supporting reports that it acted as a target regulated by SHP2 in the malignant transformation of colon cancer cells[8].The interaction probability between JAK2 and SHP2 reached $0.7586$, consistent with multiple independent studies: Ali et al. found that SHP2 participated in prolactin receptor downstream signaling by regulating SOCS-1-mediated JAK2 ubiquitination/degradation[9]; Zehender et al. further revealed that SHP2 regulated fibroblast activation by controlling TGF$\beta$-induced STAT3 signaling[10]. PTK2B and PTK2 obtained predicted probabilities of $0.7448$ and $0.7374$, respectively. Chauhan et al. confirmed that SHP2-mediated anti-apoptotic protection of multiple myeloma cells by IL-6 depends on FAK2[11]. Khare et al. reported that SHP2 differentially regulated the phosphorylation status of FAK1[12]. In the MAPK cascade, the predicted probability for MAPK14 was $0.7457$, and Chong et al. found that SHP2 promoted neuronal survival by regulating p38 and caspase activities[13]. The predicted probability for ASK1 was $0.6589$, and Yu et al. first identified SHP2 as the specific phosphatase for ASK1 Tyr-718. The study indicated that tumor necrosis factor (TNF) induced the binding of SHP2 to ASK1[14,15].

In the signal transduction/transcription factor family, the predicted probability for GRB2 was $0.7478$, indicating a strong interaction tendency between it and SHP2. Ahmed et al. demonstrated that GRB2 regulated FGFR2 phosphorylation by inhibiting receptor kinase and SHP2 phosphatase activity[16], which was highly consistent with our prediction of moderate-to-high interaction probability. The predicted probabilities for adaptor proteins GAB1 and GAB2 were $0.4767$ and $0.4425$, respectively. The classic study by Cunnick et al. elucidated that phosphorylated Tyr627 and Tyr659 on GAB1 constitute a bisphosphoryl tyrosine-based activation motif (BTAM) that mediated SHP2 binding and activation[17]. Arnaud et al. found that the SHP2-GAB2 interaction regulated IL-2-induced Rho-dependent c-fos serum response element activation[18]. These studies revealed both structurally and functionally that the Gab family constituted a classic SHP2 scaffold-type interaction partner, consistent with the interaction probability predictions for GAB1/2 in our study. The predicted probability for IRS1 is $0.5934$. Multiple studies have confirmed that SHP2/PTP2C participates in the dephosphorylation and inactivation of IRS1[19,20], indicating that the model's probabilistic prediction of IRS1-SHP2 interaction had solid biochemical support.

Among Ras pathway regulatory auxiliary proteins, in the cell adhesion/signal anchoring protein subclass, the predicted probabilities for STAT1 and STAT3 were $0.7168$ and $0.7163$, respectively. The pioneering work by Wu et al. revealed that SHP2, acting as a bispecific phosphatase, simultaneously dephosphorylates both tyrosine and serine residues of STAT1 within the cell nucleus[21]. Wang et al. demonstrated that the DDR1/SHP2 signaling complex inhibited $\alpha 2\beta 1$ integrin-mediated STAT1/3 activation[22]. The above studies collectively demonstrated a mechanism by which SHP2 regulated STAT1/3 signaling, which was highly consistent with our prediction between STAT1/3 and SHP2. The predicted probability for $\alpha$-Catenin was $0.5629$. Burks and Agazie discovered that SHP2 positively regulated FGFR3-induced cell transformation by modulating tyrosine phosphorylation of $\alpha$-Catenin[23], supporting that the interaction probability reflected SHP2's crucial regulatory role in cell adhesion-related signaling. PZR obtained the lowest predicted probability ($0.3047$). Although the study by Zhao confirmed direct binding between SHP2 and the immunoreceptor tyrosine-based inhibitory motif (ITIM) of PZR, this relatively low predicted value may reflect differences in binding modes between ITIM-mediated interactions and typical phosphatase-substrate interactions[24], this relatively low predicted value may reflect differences in the representation of ITIM-mediated interactions within our feature space compared to typical tyrosine phosphatase-substrate interactions. Consequently, the model assigned a relatively conservative score. Among pathway negative regulators/ubiquitinating enzyme subclasses, GRIN2B had a predicted probability of 0.482. Ryu et al. reviewed the functional link between SHP2 and glutamate receptors[25], suggesting that such neurotransmitter receptors may form regulatory networks with SHP2 through indirect or context-dependent mechanisms, consistent with the interaction probability provided by the model.

### S4 Supplementary Propositions and Proofs

This section establishes two properties of the distance-aware continuous-time dynamics. First, the node-specific driving term is deterministically bounded for any fixed set of trained parameters. Second, the node-state trajectory depends continuously on the normalized radial distance and is differentiable with respect to this spatial variable almost everywhere.

**Proposition 1** (Boundedness of driving terms). *For a fixed set of trained model parameters, the* *node-specific driving term*

$$f_i^{(n)} = \mathbf{W}_{\text{dyn}} g_i^{(n)} + \mathbf{b}_{\text{dyn}}, \tag{S1}$$

*is uniformly bounded over all residue nodes $i$ and continuous-time steps$n$. Specifically, there exists a finite constant $F_{\max}$ such that*

$$\left\| f_i^{(n)} \right\|_\infty \leq F_{\max}. \tag{S2}$$

*Proof.* The driving term is generated through

$$h_i^{\text{fused}} = \mathbf{W}_{\text{fuse}} \left[ h_i^{\text{agg}} \parallel e_i^{\text{dist}} \right] + \mathbf{b}_{\text{fuse}}, \tag{S3}$$

$$g_i^{(n)} = \sigma \left( h_i^{\text{fused}} \right) \in (0,1)^d, \tag{S4}$$

$$f_i^{(n)} = \mathbf{W}_{\text{dyn}} g_i^{(n)} + \mathbf{b}_{\text{dyn}}. \tag{S5}$$

Let $f_{i,\ell}^{(n)}$ denote the $\ell$th component of the driving term. It can be written as

$$f_{i,\ell}^{(n)} = \sum_{j=1}^{d} W_{\text{dyn},\ell j} g_{i,j}^{(n)} + b_{\text{dyn},\ell}. \tag{S6}$$

Because $0 < g_{i,j}^{(n)} < 1$, a deterministic lower and upper bound for each output channel can be obtained directly from the trained parameters:

$$\underline{f}_\ell = b_{\text{dyn},\ell} + \sum_{j: W_{\text{dyn},\ell j} < 0} W_{\text{dyn},\ell j}, \tag{S7}$$

$$\overline{f}_\ell = b_{\text{dyn},\ell} + \sum_{j: W_{\text{dyn},\ell j} > 0} W_{\text{dyn},\ell j}. \tag{S8}$$

Therefore,

$$\underline{f}_\ell \leq f_{i,\ell}^{(n)} \leq \overline{f}_\ell \tag{S9}$$

for every residue node $i$ and step $n$. Define

$$F_{\max} = \max_\ell \left\{ \left| \underline{f}_\ell \right|, \left| \overline{f}_\ell \right| \right\}. \tag{S10}$$

Since $\mathbf{W}_{\text{dyn}}$ and $\mathbf{b}_{\text{dyn}}$ are finite for a fixed trained model, $F_{\max} < \infty$. Consequently,

$$\left\| f_i^{(n)} \right\|_\infty \leq F_{\max}, \tag{S11}$$

which proves the result. □

Furthermore, we establish the continuous sensitivity of the entire ODE trajectory to residue distance:

**Proposition 2** (Continuous dependence and almost-everywhere sensitivity of solution trajectories to spatial position)**.** *Consider the node-level dynamics*

$$\frac{dh_i(t)}{dt} = \mathcal{F}_i \left( h_i(t), r_i \right), \tag{S12}$$

*where*

$$\mathcal{F}_i(h, r) = -\left( \boldsymbol{\tau}^{-1} + f_i(h, r) \right) \odot h + A \odot f_i(h, r). \tag{S13}$$

*Assume that $f_i(h, r)$ is continuous in $(h, r)$ and locally Lipschitz continuous in $h$. Then, on every finite interval for which the solution exists, the trajectory $h_i(t; r)$ depends continuously on the normalized radial distance $r$.*

*If $f_i(h, r)$ is differentiable at $(h_i(t; r), r)$, then the trajectory is differentiable with respect to $r$ at that point. Its sensitivity*

$$s_i(t) = \frac{\partial h_i(t; r)}{\partial r} \tag{S14}$$

*satisfies*

$$\frac{ds_i(t)}{dt} = J_i(t)s_i(t) + (A - h_i(t)) \odot \frac{\partial f_i}{\partial r}\left(h_i(t), r\right), \tag{S15}$$

*where $J_i(t) = \frac{\partial \mathcal{F}_i}{\partial h}\left(h_i(t), r\right)$ is the state Jacobian.*

*If the initial state $h_i(0) = h_{i,0}$ is independent of $r$, then*

$$\frac{\partial h_i(t; r)}{\partial r} = \int_0^t \mathbf{\Phi}_i(t, \xi) \left[(A - h_i(\xi)) \odot \frac{\partial f_i}{\partial r}\left(h_i(\xi), r\right)\right] d\xi, \tag{S16}$$

*where $\mathbf{\Phi}_i(t, \xi)$ is the state-transition matrix associated with $J_i(t)$.*

*Proof.* The node dynamics can be rewritten as

$$\mathcal{F}_i(h, r) = -\boldsymbol{\tau}^{-1} \odot h + (A - h) \odot f_i(h, r). \tag{S17}$$

Because $f_i(h, r)$ is continuous in $(h, r)$ and locally Lipschitz in $h$, the vector field $\mathcal{F}_i(h, r)$ has the same local continuity and Lipschitz properties. Standard parameter-dependence results for ordinary differential equations therefore imply that the solution $h_i(t; r)$ depends continuously on $r$ on every finite interval over which the solution exists.

At points where $f_i$ is differentiable with respect to $h$ and $r$, define

$$s_i(t) = \frac{\partial h_i(t; r)}{\partial r}. \tag{S18}$$

Differentiating

$$\frac{dh_i(t)}{dt} = \mathcal{F}_i\left(h_i(t), r\right) \tag{S19}$$

with respect to $r$ and applying the chain rule gives

$$\frac{ds_i(t)}{dt} = \frac{\partial \mathcal{F}_i}{\partial h}\left(h_i(t), r\right) s_i(t) + \frac{\partial \mathcal{F}_i}{\partial r}\left(h_i(t), r\right). \tag{S20}$$

The direct derivative of the vector field with respect to $r$, with $h$ held fixed, is

$$\begin{aligned} \frac{\partial \mathcal{F}_i}{\partial r} &= -\frac{\partial f_i}{\partial r} \odot h_i + A \odot \frac{\partial f_i}{\partial r} \\ &= (A - h_i) \odot \frac{\partial f_i}{\partial r}. \end{aligned} \tag{S21}$$

Let

$$J_i(t) = \frac{\partial \mathcal{F}_i}{\partial h}\left(h_i(t), r\right) \tag{S22}$$

and

$$b_i(t) = (A - h_i(t)) \odot \frac{\partial f_i}{\partial r}\left(h_i(t), r\right). \tag{S23}$$

The sensitivity equation can then be expressed as the linear non-homogeneous system

$$\frac{ds_i(t)}{dt} = J_i(t)s_i(t) + b_i(t). \tag{S24}$$

Let $\mathbf{\Phi}_i(t,\xi)$ denote the state-transition matrix of the homogeneous system

$$\frac{dz(t)}{dt} = J_i(t)z(t), \tag{S25}$$

with

$$\mathbf{\Phi}_i(\xi,\xi) = I. \tag{S26}$$

By the variation-of-constants formula,

$$s_i(t) = \mathbf{\Phi}_i(t,0)s_i(0) + \int_0^t \mathbf{\Phi}_i(t,\xi)b_i(\xi)\,d\xi. \tag{S27}$$

Because the initial representation is assumed to be independent of the radial distance,

$$s_i(0) = \frac{\partial h_{i,0}}{\partial r} = 0. \tag{S28}$$

Substitution of $b_i(\xi)$ therefore yields

$$\frac{\partial h_i(t;r)}{\partial r} = \int_0^t \mathbf{\Phi}_i(t,\xi)\left[(A - h_i(\xi)) \odot \frac{\partial f_i}{\partial r}(h_i(\xi),r)\right] d\xi. \tag{S29}$$

□

Together, these propositions establish that the node-specific driving terms of LNGCN are bounded and provide a mathematically explicit pathway through which normalized radial distance can influence residue-state trajectories. They demonstrate the capacity of the continuous-time dynamics to incorporate residue-specific spatial information, but do not imply that every pair of distinct radial positions must produce distinct representations. The practical preservation of residue-level representational heterogeneity was therefore evaluated separately using node variance, Dirichlet energy, pairwise representation distance, radial-bin separation, and effective embedding rank.

### S5 Calibration method

The Platt scaling calibration method[26,27] is a classic parametric probability calibration approach that maps raw scores to approximate posterior probabilities by fitting a one-dimensional logistic regression model to the logits produced by the model. Let the uncalibrated logit of a protein for a sample be denoted as $z \in \mathbb{R}$. The Platt scaling formula is:

$$\hat{p}(y = 1 \mid z) = \sigma(Az + B) = \frac{1}{1 + e^{-(Az+B)}}, \tag{S30}$$

where $A$ and $B$ are parameters fitted on the calibration data, and $\sigma(\cdot)$ denotes the sigmoid function. This method preserves the ranking of samples while correcting the issue of overall confidence being systematically too high or too low.

The core idea of Beta calibration[28,29] is to fit logistic regression in a log-probability space. This method can better describe skewed or heavy-tailed probability distributions. For a given logit $z$, the uncalibrated probability is first computed as:

$$p = \sigma(z) = \frac{1}{1 + e^{-z}}. \tag{S31}$$

To avoid numerical issues, $p$ is clipped to $[\varepsilon, 1 - \varepsilon]$, where $\varepsilon = 10^{-7}$. Subsequently, a two-dimensional feature is constructed: $x = [\log p,\ \log(1 - p)]$ and logistic regression is fitted in this feature space:

$$\hat{p}(y = 1 \mid p) = \sigma(a \log p + b \log(1 - p) + c), \tag{S32}$$

Isotonic calibration[30,31] is a non-parametric, monotonic-constrained probabilistic calibration method. Its objective is to learn a monotonic non-decreasing function $f$ over the interval $[0, 1]$, such that for any observation pair $(p_i, y_i)$ consisting of an initial uncalibrated probability $p$ and a label $y \in \{0, 1\}$, the following holds:

$$\hat{p}(y = 1 \mid p) = f(p), \quad f \text{ is non-decreasing}, \tag{S33}$$

and typically estimates $f$ by minimizing the squared loss $\sum_i (f(p_i) - y_i)^2$. Unlike the aforementioned parameterized methods, Isotonic regression does not presuppose the functional form, requiring only monotonicity, thus exhibiting strong fitting capabilities when training data is abundant.

The Boosting calibration[31,32] can further capture potentially highly nonlinear, even locally non-monotonic calibration relationships. In this study, we introduced a Gradient Boosting-based calibrator, whose approach involved directly using the logit $z$ as a one-dimensional input feature to train a gradient boosting binary classification model $g(z)$:

$$\hat{p}(y = 1 \mid z) = g(z), \tag{S34}$$

where $g(\cdot)$ is composed of multiple decision trees added together, with each tree sequentially fitting the residuals from the previous round to minimize the logistic loss.

### S6 Supplementary experimental methods

#### S6.1 Complete medium compositions

HeLa cells (European Collection of Authenticated Cell Cultures) were maintained in DMEM (Sigma-Aldrich, #D5671) supplemented with: 10% FBS (Vistech, #SE100-B), 1 mM sodium pyruvate (Gibco, #11360070), $1\times$ MEM Non-Essential Amino Acids (Gibco, #11140050), 2 mM GlutaMAX (Gibco, #35050061), 100 U/mL penicillin and 100 $\mu$g/mL streptomycin (Gibco, #15140122).

#### S6.2 Detailed DNA and siRNA Transfection protocols

For DNA transfections, six million HeLa cells were seeded on eight 145-mm dishes (four dishes per condition) and transfected the next day with the vector expressing TPR (pcDNA4/TO-TPR-2siR-EGFP) or the empty vector (pcDNA4/TO-EGFP) as a negative control, using Neofect DNA Transfection Reagent (Neofect, #TF20121201) according to the manufacturer's protocol. 24 hours after transfection, the medium was replaced, and cells were harvested for immunoprecipitation 48 hours after transfection.

For siRNA transfections, six hundred thousand HeLa cells were seeded on 6-cm dishes and transfected the next day using Lipomaster 3000 Transfection Reagent (Vazyme, #TL301-02) according to the manufacturer's instructions. siRNAs were used at a final concentration of 25 nM. Twenty-four hours after transfection, cells were passaged at a 1:3 ratio and harvested for immunoblotting 72 hours after transfection. The specific oligonucleotides used for each assay in this study are listed in Table S4.

Table S4: siRNA sequence

| Sequence | Name | siRNA | Source |
|---|---|---|---|
| 5'-GAUCAAGUCCAAUAUCGAUtt-3' | siRALY | siRNA against RALY | Tsingke |
| 5'-AUCGAUAUUGGACUUGAUCtg-3' | siRALY | siRNA against RALY | Tsingke |
| 5'-ACUUAUUCGGGAUAAAGUAtt-3' | siELAVL1 $\#1$ | siRNA against ELAVL1 $\#1$ | Tsingke |
| 5'-UACUUUAUCCCGAAUAAGUtt-3' | siELAVL1 $\#1$ | siRNA against ELAVL1 $\#1$ | Tsingke |
| 5'-UGAACUACGUGACCGCGAAtt-3' | siELAVL1 $\#2$ | siRNA against ELAVL1 $\#2$ | Tsingke |
| 5'-UUCGCGGUCACGUAGUUCAca-3' | siELAVL1 $\#2$ | siRNA against ELAVL1 $\#2$ | Tsingke |

#### S6.3 Immunoprecipitation Buffer Details

The composition of lysis buffer is 20 mM Tris (pH 8.0), 150 mM NaCl, 0.5% NP-40, 1 mM EDTA, 1 mM $MgCl_2$, protease inhibitors (Yeasen, #20124ES), phosphatase inhibitors (Yeasen, #20109ES), benzonase nuclease (Merck, #70664, 50 U/mL final concentration). And dilution buffer has the same composition as the lysis buffer without benzonase. As for SDS sample buffer, it contains 2% SDS, 10% glycerol, 50 mM Tris-HCl, pH 6.8.

#### S6.4 Antibody Information

Primary antibodies: ELAVL1 (Proteintech, #11910-1-AP, rabbit, 1:2000), GFP (Huabio, #ET1607-31, rabbit, 1:2500), RALY (Proteintech, #68011-4-Ig, mouse, 1:1000), $\beta$-tubulin (Huabio, #ET1602-4, rabbit, 1:20000). Secondary antibodies: goat anti-mouse IgG H&L HRP (Abcam, #ab205719, 1:5000), goat anti-rabbit IgG H&L HRP (Abcam, #ab205718, 1:5000). And SDS sample buffer contains 2% SDS, 10% glycerol, 50 mM Tris-HCl, pH 6.8.
